# Glycoengineering with neuraminic acid analogs to label lipooligosaccharides and detect native sialyltransferase activity in Gram-negative bacteria

**DOI:** 10.1101/2024.07.05.602201

**Authors:** Erianna I. Alvarado Melendez, Hanna de Jong, Jet E. M. Hartman, Jun Yang Ong, Marc M. S. M. Wösten, Tom Wennekes

## Abstract

Lipooligosaccharides (LOS) are the most abundant cell surface glycoconjugates on the outer membrane of Gram-negative bacteria. They play important roles in host-microbe interactions. Certain Gram-negative pathogenic bacteria cap their LOS with the sialic acid, *N*-acetylneuraminic acid (Neu5Ac), to mimic host glycans that among others protects these bacteria from recognition by the hosts immune system. This process of molecular mimicry is not fully understood and remains under investigated. To explore the functional role of sialic acid-capped lipooligosaccharides (LOS) at the molecular level, it is important to have tools readily available for the detection and manipulation of both Neu5Ac on glycoconjugates and the involved sialyltransferases, preferably in live bacteria. We and others have shown that the native sialyltransferases of some Gram-negative bacteria can incorporate extracellular unnatural sialic acid nucleotides onto their LOS. We here report on the expanded use of native bacterial sialyltransferases to incorporate neuraminic acids analogs with a reporter group into the LOS of a variety of Gram-negative bacteria. We show that this approach offers a quick strategy to screen bacteria for the expression of functional sialyltransferases and the ability to use exogenous CMP-Neu5Ac to decorate their glycoconjugates. For selected bacteria we also show this strategy complements two other glycoengineering techniques, Metabolic Oligosaccharide Engineering (MOE) and Selective Exo-Enzymatic Labelling (SEEL), and that together they provide tools to modify, label, detect and visualize sialylation of bacterial LOS.

## Introduction

Cell surface glycoconjugates play a key role in mediating interactions between bacteria and the host (Poole et al., 2018). One important and prevalent monosaccharide found at the terminal end of several classes of host glycoconjugates is *N*-acetylneuraminic acid (Neu5Ac), commonly known as sialic acid. In mammals, it is found in gangliosides, *N*-/*O*-linked glycoproteins, and polysialic acid (Lewis et al., 2015), where it has important functional roles such as mediation of cell-cell interactions, modulation of the immune response, cell signaling (Chang and Nizet, 2014; Varki and Gagneux, 2012), but also serves as a binding site for pathogens (Hirmo et al., 1998).

Compared to mammals, Neu5Ac is less commonly found in Gram-negative bacteria. Currently, a limited number of bacterial species are known to display Neu5Ac as part of their capsular polysaccharides (CPS), or their lipopolysaccharides (LPS) and lipooligosaccharide (LOS). Some of these species can produce Neu5Ac *de novo*, and others have to scavenge it from the host such as *Neisseria gonorrhoeae* (Ghosh, 2020). In particular, sialylated LOS has been found in strains of *Campylobacter jejuni, Vibrio, Neisseria*, and *Haemophilus influenza* (Dudek et al., 2022). Sialylation of bacterial LOS occurs via sialyltransferases (ST) and can result in the mimicry of host glycans, a phenomenon observed in some pathogens such as *N. gonorrhoeae, H. influenzae* and *C. jejuni* (Moran et al., 1996).

Bacteria can use this strategy of mimicking host glycans to mask their surface and avoid being recognized as foreign cells by the immune system. There are several mechanisms by which host cells distinguish between self and non-self cells. One such mechanism is the alternative complement pathway. This pathway uses factor H, surface-bound C3b and host polyanions to discriminate between cells (Pangburn et al., 2008). It is known that cells containing sialic acid do not activate this pathway (Pangburn et al., 2008). Another way of distinguishing host from foreign cells is through sialic acid binding receptors (Siglecs). Siglecs recognize sialic acid-containing ligands expressed on the surface of cells, the presence of these ligands attenuates inflammatory and innate immune responses of Toll-like receptors (Crocker et al., 2007; Paulson et al., 2012; Varki and Gagneux, 2012). In this way, bacterial glycans containing Neu5Ac can be recognized by host Siglecs and factor H, which leads to dampening of immune signals or prevents the activation of the alternative complement pathway, respectively (de Jong et al., 2022a; Kopp et al., 2012; Leaubli et al., 2022; Meri and Pangburn, 1990). Mimicry through sialic acid can also induce autoimmune mechanisms leading to the development of diseases like Guillain-Barré syndrome, which is caused by certain strains of *C. jejuni*. In the case of pathogenic Neisseria strains, sialylation of the LOS has been reported to influence the susceptibility of the bacterium to bactericidal antibodies (Rest and Frangipane, 1992). This sialylation has a negative impact on the host complement pathway activation, preventing phagocytosis, reducing adhesion to neutrophils and thus also induction of an oxidative burst (Carlin et al., 2009; Moran et al., 1996; Rest and Frangipane, 1992; Wetzler et al., 1992). For these reasons, the presence of Neu5Ac in bacterial LOS is considered a virulence factor, yet the precise functional role of Neu5Ac in molecular mimicry is not fully understood yet. Additionally, there may be other bacteria capable of decorating their LOS with sialic acid that have yet to be discovered. To investigate this and gain deeper insights, we need glycoengineering techniques to detect strains with active sialyltransferases and manipulate Neu5Ac in bacterial glycoconjugates.

Currently, there are several widely applied approaches to perturb and study the role of glycans in mammalian cells (Critcher et al., 2021; Edgar, 2021; Griffin and Hsieh-Wilson, 2016; Nischan and Kohler, 2016), but only a few techniques are available to study the bacterial glycome (Calles-Garcia and Dube, 2024). The most widely used approach is Metabolic Oligosaccharide Engineering (MOE). In this technique carbohydrates modified with bioorthogonal reporters are taken up by the cell, metabolically processed and transferred onto the cellular glycans. The metabolically incorporated glycan analog can be studied via its reporter by covalently attaching a fluorophore or an enrichment tag. MOE has been applied to microbes to engineer their cell wall (Dumont et al., 2012; Liu et al., 2009; Sadamoto et al., 2004, 2002; Swarts et al., 2012), image (Geva-Zatorsky et al., 2015; Hudak et al., 2017), discover glycoproteins (Besanceney-Webler et al., 2011; Champasa et al., 2013) or develop new antibacterial strategies (Kaewsapsak et al., 2013; Memmel et al., 2013; Tra and Dube, 2014). Although MOE is a very useful technique, it has limitations. To modify glycans, the carbohydrate analogs used must be able to enter the cell, be metabolically processed by the native salvage pathway enzymes and finally be accepted as substrates by native intracellular glycosyltransferases. Thus, this technique cannot be applied to bacteria that lack the biosynthetic machinery to metabolically process the monosaccharide. Furthermore, shortcomings in any of these steps often result in limited control over the level and location of reporter incorporation, *e*.*g*. glycoproteins or glycolipids (Bussink et al., 2007). The metabolically generated sugar nucleotide analogs are often substrates for several related intracellular glycosyltransferases. Therefore, MOE may also result in different linkage types, e.g. α2,3-vs. α2,6-sialic acid linkages, which complicate their subsequent analysis and interpretation of their biological effects. In addition, the efficacy and precision of this technique are significantly influenced by the sugar analog employed. For instance, in mammals, UDP-GalNAc and UDP-GlcNAc can be interconverted by epimerases, which in turn are utilized by different glycosyltransferases, resulting in the incorporation of these probes on distinct glycans (Boyce et al., 2011). Another disadvantage is the toxicity associated with carbohydrate analogs containing azido moieties and Cu(I), when copper-catalyzed azide-alkyne cycloaddition (CuAAC) is used (Han et al., 2018; Liu et al., 2022; Saïdi et al., 2022).

Another glycoengineering technique is selective exoenzymatic labelling (SEEL) (Sun et al., 2016; Yu et al., 2016) that we have applied to label the LOS of *N. gonorrhoeae* (de Jong et al., 2022b). SEEL uses an externally applied recombinant glycosyltransferase to selectively label the outside of a cell with tailor-made sugar nucleotide analogs. The advantages of SEEL are the precise incorporation of a monosaccharide derivative onto a defined acceptor on the cell surface with a known linkage type, and low cytotoxicity. The established linkage type can also be non-native to the bacterium, such as α-2,6-Neu5Ac instead of the naturally occurring α-2,3-Neu5Ac on *N. gonorrhoeae* (Gilbert et al., 1996), which is a unique feature of SEEL compared to other glycoengineering techniques. One disadvantage of this technique is the need for a specific recombinant sialyltransferase that can recognize the LOS epitope in the bacterium of interest. For several bacteria the LOS structures are still not known. When applying SEEL to *N. gonorrhoeae* (de Jong et al., 2022b), we observed that wild-type bacteria were able to incorporate the sugar nucleotide analogs with their own sialyltransferases. *N. gonorrhoeae* was previously reported to scavenge sialic acid nucleotides from the environment and transfer them to its lipooligosaccharides via native sialyltransferases (Gulati et al., 2020, 2015; Jen et al., 2021; Nairn et al., 1988). In our experiments, we also noticed that the level of sialylation achieved when using SEEL in *N. gonorrhoeae* was lower than the produced by the bacterium native sialyltransferases (de Jong et al., 2022b).

Intrigued by the possibility of using endogenous enzymes to engineer the cell surface glycoconjugates of bacteria, we sought to further investigate this labelling strategy on two other pathogens known for displaying sialylated LOS. We selected Non-typeable *H. influenzae* (NTHi), and *C. jejuni* (strains associated with Guillain-Barré), to assess the labeling of their own LOS using their native sialyltransferases (NSTs) and CMP-Neu5Ac analogs and in addition performed a qualitative comparison between this approach, MOE and SEEL. To further extend the application of glycoengineering via NSTs to more bacteria, we performed a BLAST search using as input the annotated sialyltransferases found in *N. gonorrhoeae* (*lst*), *C. jejuni* (*CstII*) and NTHi (*lic3A, lsgB* and *SiaA*). From the BLAST search, we selected 49 strains of Gram-negative bacteria that includes relevant human pathogens, commensal, and zoonotic species. We applied our labelling strategy to label the lipooligosaccharides and detect native sialyltransferase activity in this selection of Gram-negative bacteria with Neu5Ac analogs. This revealed that a variety of pathogenic and commensal bacteria can utilize CMP-Neu5Ac analogs to label their LOS. For some of the bacteria, this represents the first reported evidence of sialyltransferase activity. It also showcased the ease by which bacterial native sialyltransferase activity and sialylated surface glycans can be detected via this approach.

## Results and discussion

### Comparing bacterial glycoengineering via NSTs, MOE and SEEL

For its glycan mimicry (de Jong et al., 2022b), *N. gonorrhoeae* transfers scavenged CMP-Neu5Ac onto its LOS with an α-2,3 linkage with its sialyltransferase *lst* (Mubaiwa et al., 2017). As *N. gonorrhoeae* lacks the genes for the biosynthesis of CMP-Neu5Ac, MOE is not possible (Figure 1A). This was verified by probing both the wild type *N. gonorrhoeae* and a sialyltransferase mutant (Δ*lst*) with known MOE probe, Neu5Az, followed by a Strain-Promoted Alkyne-Azide Cycloaddition reaction (SPAAC) click reaction and observing no labelling of the LOS (Figure 1A). We previously showed that *N. gonorrhoeae* LOS can be labelled using SEEL (de Jong et al., 2022d). We also found that endogenous sialyltransferases are able to transfer CMP-Neu5Az onto their LOS, something that has also been observed by other groups. (Gulati et al., 2020, 2015; Jen et al., 2021), We first wanted to further investigate the substrate scope of the *lst* native sialyltransferase and the speed of the NST approach. For this we exposed *N. gonorrhoeae* to two different analogs of the sugar nucleotide donor: the known CMP-Neu5Az (Figure 1B), and the not yet tested CMP-Neu5biotin (Figure 1C) and observed that LOS labelling using CMP-Neu5Az occurs quickly, within minutes, after incubation with the probe (Figure 1D). Unsurprisingly, we also found that CMP-Neu5biotin was incorporated in the LOS, showing that larger modifications at the fifth position are accepted by *lst*, as was already observed in our previous work when using CMP-Neu5AF_488_ (de Jong et al., 2022b).

**Figure 1.**
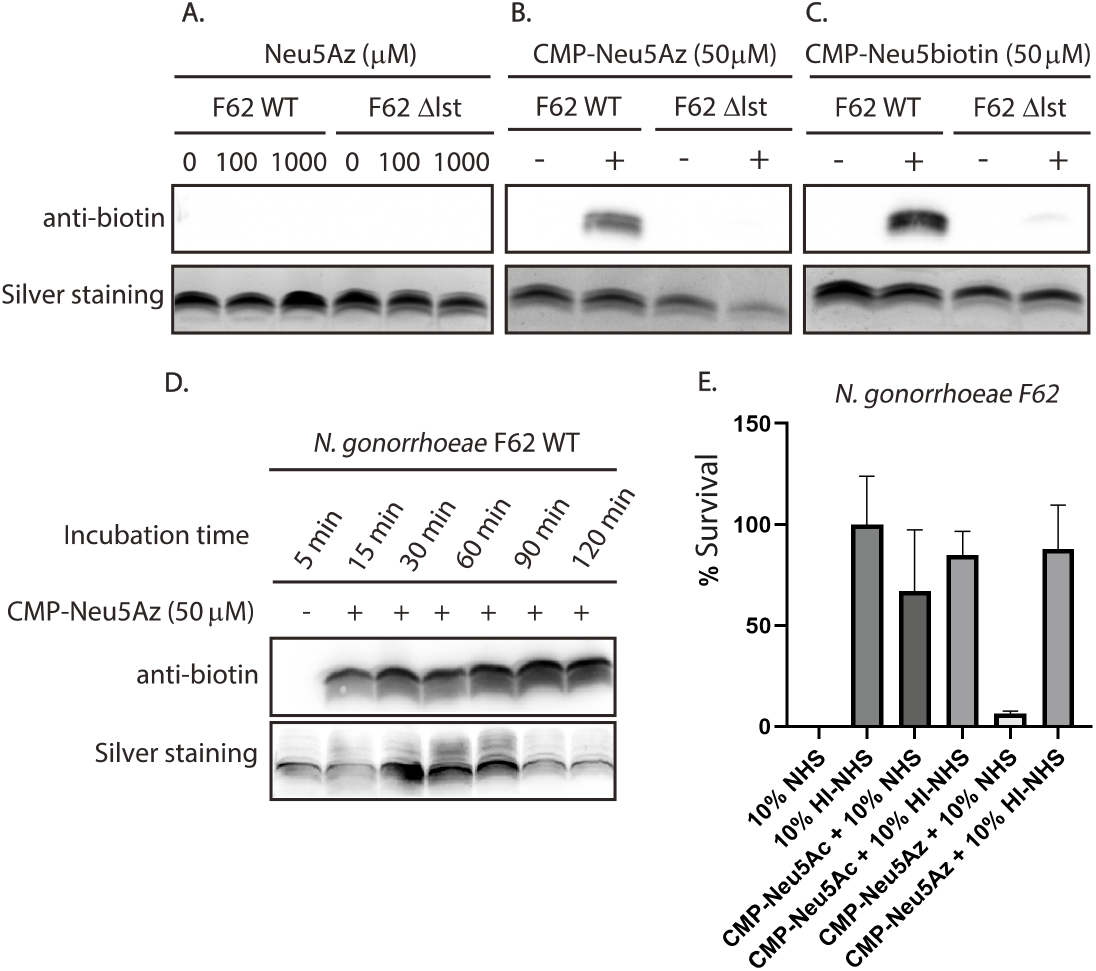
MOE and NSTs applied to *Neisseria gonorrhoeae*. (A) MOE with Neu5Az on *N. gonorrhoeae* wild type and the ST mutant (Δ*lst*) followed by a SPAAC with DBCO-PEG_4_-biotin. (B) Labelling of *N. gonorrhoeae* WT LOS by NSTs followed by SPAAC using DBCO-PEG_4_-biotin. (C) Incorporation of Neu5biotin into the Neisserial LOS by NSTs. LOS was analyzed on Western blot and by silver staining. (D) Incubation of *N. gonorrhoeae* with CMP-Neu5Az at different time points followed by SPAAC click with DBCO-PEG_4_-biotin. (E) Serum resistance of *N. gonorrhoeae* which was grown without or with a nucleotide sugar; CMP-Neu5Ac or CMP-Neu5Az. Colony forming units were counted after incubation in the presence of Normal Human Serum (NHS) or Heat inactivated NHS (HI-NHS), and reported as % survival.

Additionally, to show the utility of this labelling method to study the effect of bacterial sialylation on serum-mediated killing we performed an assay using the CMP-Neu5Az. Gulati et al., previously showed that *N. gonorrhoeae* has altered resistance against serum-mediated killing when the bacteria were modified with other sialic acids than Neu5Ac, such as legionaminic acid (Gulati et al., 2020, 2015). However, sialylation with Neu5Az has not been reported yet. In a serum-resistance assay with bacteria covered by Neu5Az, the bacteria were less resistant against serum compared to bacteria treated with native CMP-Neu5Ac (Figure 1E).This observation fits the trend that a small modification on the glycan can perturb the interaction with factors from serum, which also has been seen for the 9-azido derivative of the nucleotide sugar (Gulati et al., 2015).

The next Gram-negative bacterium we investigated, non-typeable *H. influenzae* (NTHi), is known for displaying sialic acid on its LOS to increase resistance to serum-mediated killing (Apicella, 2012; de Jong et al., 2022a; Heise et al., 2018). We started our investigation by corroborating the results obtained by Heise et al. when applying MOE to NTHi (Heise et al., 2018). We found that Neu5Az was incorporated into its LOS and not by a mutant lacking the sialic acid transporter *SiaP* (Figure 2A). We did not observe incorporation of Neu5Az into other cell surface glycoproteins (Supporting information, Figure S1), but did notice a decrease in bacterial growth, indicative of cell toxicity, when using a high concentration of Neu5Az (1 mM) (Supporting information, Figure S2).

**Figure 2.**
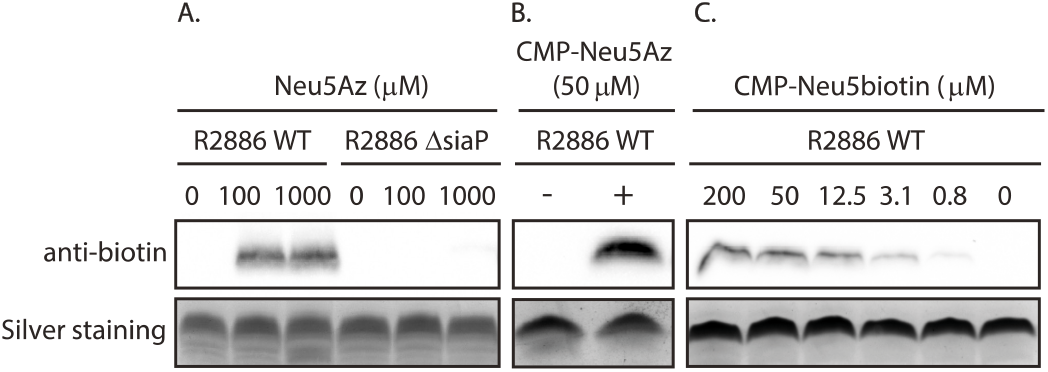
(A) MOE applied to label LOS of NTHi wildtype (WT), and a sialic acid transporter mutant (Δ*siaP*) using Neu5Az followed by SPAAC reaction with DBCO-PEG4-biotin. (B) Labelling of LOS by NSTs of NTHi using CMP-Neu5Az, followed by SPAAC with DBCO-PEG_4_-biotin. (C) Labelling of NTHi LOS by the NSTs using a concentration range of CMP-Neu5biotin.

We next tested SEEL to label NTHi using CMP-Neu5biotin in combination with sialyltransferases of mammalian origin first (ST6Gal1) and then STs from bacteria (Pmst1, Pmst3 and Pd2,3). We did not observe any labelling of NTHi LOS, with the different enzymes tested, which indicated that to achieve labeling via the SEEL technique, the selection of STs is essential, and the method is compromised for bacteria in which the LOS acceptor structure or LOS shielding is unknown (Supporting information, Figure S3). We however did observe that in the absence of exogenous enzyme, the NSTs from NTHi can incorporate the unnatural Neu5Ac analog into the LOS (Figure 2B). We tested the ability of the native sialyltransferases using CMP-Neu5Az, and CMP-Neu5biotin, at different concentrations. With both nucleotide sugars we observed labelling of the LOS. When using CMP-Neu5biotin we observed labelling up to a concentration of as low as 0.8 μM (Figure 2C).

We continued our investigation with *C. jejuni*, the major cause of bacterial gastroenteritis in the developing world. Certain *C. jejuni* strains are also linked to the autoimmune disease Guillain-Barré syndrome (GBS) and have unique LOS structures that mimic the terminal epitopes of human glycans. *C. jejuni*’s LOS can be sialylated and bind to host receptors, such as Siglec 1 and 7 (A. P. Heikema et al., 2013; Astrid P. Heikema et al., 2013). This mimicry can however also lead to the production of anti-ganglioside antibodies targeting the nerves of the host during the disease, which can manifest in a variety of ways, including symptoms such as mild paralysis to acute pain (van den Berg et al., 2014; Yuki et al., 2004). Due to the link between the sialylated LOS of *C. jejuni* and the development of GBS, this is an intriguing target to manipulate and study. LOS sialylation occurs via sialyltransferases *CstI* and *CstII* that create either α-2,3-or both α-2,3- and α-2,8-linkages depending on a point mutation (Gilbert et al., 2002; Godschalk et al., 2007). In our study we focused on two strains of *C. jejuni* that are associated with GBS: GB11 and GB19, along with the corresponding mutant strains of *CstII* (Godschalk et al., 2007, 2004; Louwen et al., 2008).

We first tested MOE by using Neu5Az to attempt labeling *C. jejuni*’s LOS. For the WT strains we observed labelling and as expected not for the sialyltransferase mutants (Figure 3A). This result is the first properly documented example of MOE in *C. jejuni*. Next, SEEL was investigated in the *CstII* mutants of all three strains using recombinant *CstII* as the external sialyltransferase. The ST, *CstII*, has been characterized as a bifunctional sialyltransferase having both α2,3-sialyltransferase and α2,8-sialyltransferase activities, with a preference for α2,3-sialosides substrates (Babulic et al., 2023). We found that under our experimental conditions, *CstII* can sialylate the LOS of strain GB11, but not the LOS of strain GB19 (Supporting information, Figure S4). We hypothesize that, since the LOS structure of these strains differ, strain GB19 probably requires another sialyltransferase. We therefore tested SEEL on *C. jejuni* GB19 using a combination of *CstII* and *CstI* (Chiu et al., 2004), and labelling was indeed observed (Supporting information, Figure S5). With SEEL in these GB *Campylobacter* strains proving successful, we tested the ability of the native STs in the WT strains to transfer the nucleotide sugar derivative. Figures 3B-C show successful labelling of the LOS using CMP-Neu5Az, CMP-Neu5biotin and with a fluorescent reporter group (CMP-Neu5AF_488_). No labelling was observed for the strains lacking the *CstII* genes, as we demonstrated with the *CstII* mutants (Δ*cstII*) of the GB strains 11 and 19.

**Figure 3.**
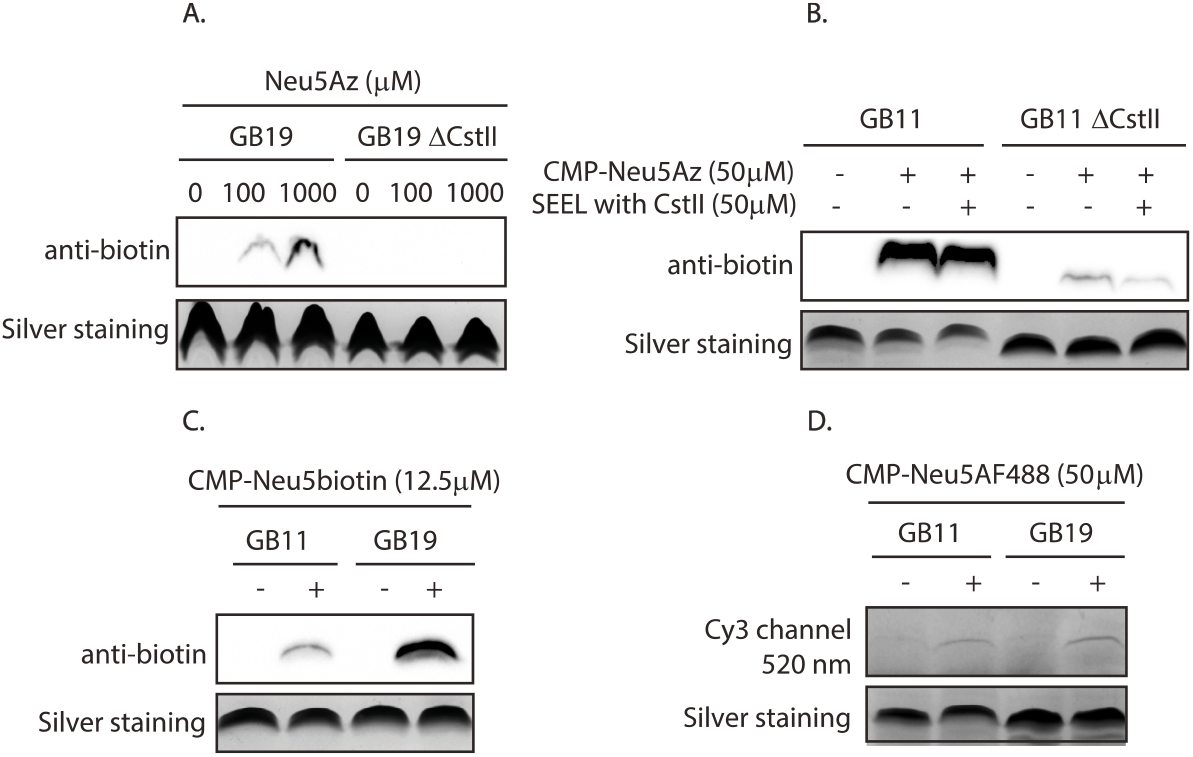
MOE, SEEL and native sialyltransferases, applied to *C. jejuni*. (A) MOE with Neu5Az on GB19 wildtype and the *CstII* sialyltransferase mutant. The probe is incorporated into the LOS of the WT, as observed after a SPAAC reaction with DBCO-PEG_4_-biotin. (B) SEEL with exogenous applied *CstII*. (C) Labelling of LOS by the native sialyltransferases using CMP-Neu5biotin. (C) In-gel fluorescence of LOS modified by native sialyltransferases with CMP-Neu5AF_488_.

After having established that strains of NTHi, *N. gonorrhoeae* and *C. jejuni* could incorporate modified nucleotide sugars via their own native sialyltransferases, we sought to investigate whether other bacterial species were also capable of this activity and to develop a strategy to use CMP-Neu5Ac analogs to scan bacteria for the expression of active STs and the capacity to use exogeneous CMP-Neu5Ac to sialylate their LOS.

### Scanning for sialyltransferase activity and sialylated LOS in other pathogenic bacteria

We started with a BLAST search on the six sialyltransferase genes known or expected to be capable of ST activity in NTHi, *N. gonorrhoeae* and *C. jejuni*: *lst* (α2,3-ST, from *N. gonorrhoeae* strain F62), *Lic3A* (α2,3-ST, from NTHi R2886), *SiaA* (α2,8-ST, from NTHi), *LsgB* from NTHi, *CstII* (α2,3-ST, *C. jejuni*) and *PmST1* (α2,3-ST from *P. multocida* pm70) (see Supporting information Table 2). From the hits of this BLAST search, we have selected 49 Gram-negative bacteria strains, among them human pathogens, commensal and zoonotic species. This list includes representatives of the Campylobacter genus, such as *C. insulaenigreae, C. upsaliensis, C. lari* and *C. coli*; and Pasteurella species such as *P. dagmatis* and *P. multocida*. We also included *Neisseria meningitidis* strains, a variety of Haemophilus strains, *Prevotella timonensis* and *Prevotella bivia* species and commensal bacteria such as *Neisseria lactamica* and *Haemophilus parainfluenzae*, to make the selection as diverse as possible.

The activity of the sialyltransferases of these bacteria was tested using our previously optimized standardized conditions, developed during the LOS labelling of NTHi, *N. gonorrhoeae* and *C. jejuni*. We however do not exclude the possibility that the sialyltransferase activity of these bacteria might alter under different labelling, environmental or culturing conditions. In all the experiments, we examined the labelling of cell surface glycoconjugates such as LPS/LOS, and also the possible incorporation of Neu5Az into cell-surface glycoproteins and capsular polysaccharides (CPS) in those bacteria that produce them.

We used a two-step labelling approach to label bacterial LOS by incubating the bacteria with CMP-Neu5Az, followed by a SPAAC reaction using DBCO-PEG_4_-biotin. For some experiments we also use compared this to CuAAC for the click reaction and obtained identical results (Supporting information, Figure S6). To visualize the potentially labeled LOS, samples were digested with Proteinase K overnight, and boiled in loading buffer before loading onto tris-tricine gels. It is worth mentioning that most of the LOS structures labelled in the experiments reported here have not yet been characterized or reported in the literature, so we could not directly relate the results to the expected size of the LOS.

We observed labeling of the LOS/LPS in 39 out of 49 selected bacteria. Among these: *Prevotella timonensis, Haemopilus aegyptius, Haemophilus parainfluenzae, Campylobacter upsaliensis, Campylobacter insulaenigrae, Pasteurella multocida, Pasteurella dagmatis, Neisseria lactamica*, and *Neisseria meningitidis*. These results indicate that these bacteria are expressing active STs capable of utilizing exogenous CMP-Neu5Az to label their LOS. However, it should be emphasized that the absence of labelling in some experiments does not exclude the presence of sialyltransferases in those bacteria.

Protein sialylation was evaluated by running samples on SDS gels without the Proteinase K digestion step. We also assessed the labelling of CPS of strains *C. jejuni* GB11 and GB19, *N. gonorrhea* and *N. meningitidis* B, C and W135; we did not observe labelling of proteins or CPS for any of the bacteria when incubated with CMP-Neu5Az (Supporting information, Figure S7 and S8).

Sialylation via the method we employ requires an active sialyltransferase, a substrate CMP-Neu5Az, and an appropriate acceptor epitope in the LOS. Our experiments represent a situation for these bacteria where they are in a simulated environment rich in their ST sugar-nucleotide substrate. The detection of sialyltransferase activity via our method hints at a mechanism by which these bacteria can use CMP-Neu5Ac available from the environment to cap their LOS/LPS. The short incubation time required for labelling, as seen in *N. gonorrhoeae* (Figure 1D), indicates that the STs could be located at or near the bacterial surface (Shell et al., 2002).

It is known that the source of Neu5Ac for *N. gonorrhoeae* originates from human erythrocytes (Nairn et al., 1988). Recent studies have shown that platelets are a source of nucleotide sugars, that can be released after a stress trigger, and that sialylation via this process is possible *in vivo* (Jones et al., 2016; Lee et al., 2014; Manhardt et al., 2017). Platelets or other sources such as liberation in co-culture with other bacteria, are a plausible source of nucleotide sugars. An alternative mechanism for the incorporation of nucleotide sugars could be the uptake of these nucleotide sugars from the environment and sialyltransferase activity in the cytoplasm instead of on the surface. From our data it is not possible to draw definite conclusions about the exact location of the sialyltransferases. We previously reported microscopy data for *N. gonorrhoeae* incubated with CMP-Neu5AF_488,_ which showed no fluorescence in the cytoplasm (de Jong et al., 2022b). Since the fluorescence signal was not distributed throughout the cytoplasm and was localized outside the membrane, this tentatively indicates that the sialyltransferases are active in the periplasm or on the surface. In addition, we here fluorescently labeled another strain, *N. meningitidis B* with CMP-Neu5AF_488_, and observed a similar localization (See Figure 4). We excluded the possibility that the hydrolysis of the sugar nucleotides provides sialic acid analogs that are then metabolized and incorporated into the LOS, since LOS labelling was observed almost immediately after incubation with CMP-Neu5Az in the case of *N. gonorrhoeae* (Figure 1D). The exact location of the sialyltransferase is an outstanding question for most of these bacteria and it is our future aim to investigate for the various bacteria used here the location of the sialyltransferases, and whether the nucleotide sugars are taken up and metabolized intracellularly or transferred via ectopic glycosyltransferases from bacteria.

**Figure 4.**
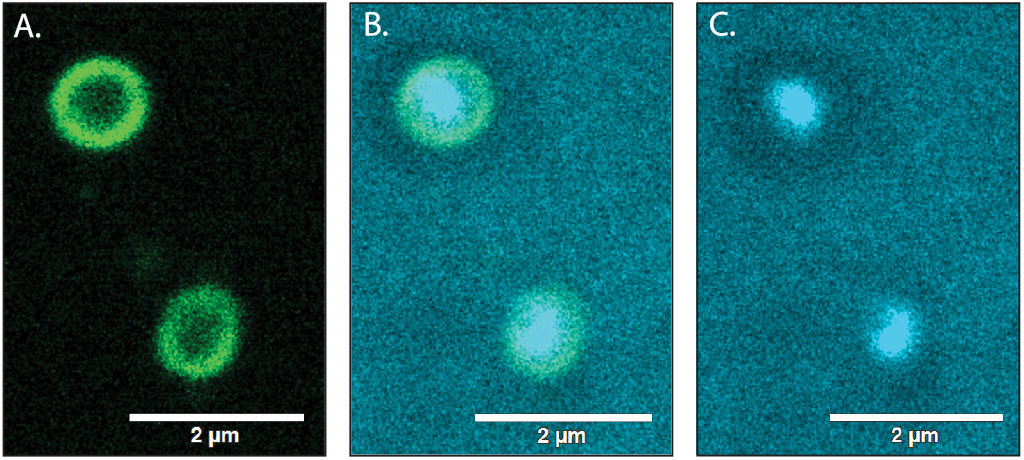
(A) Fluorescence microscopy images of *N. meningitidis* B fluorescently labeled using CMP-Neu5AF_488_ (488 channel). (B) Fluorescence microscopy images of DAPI stain 405 channel (C) Merge of 488 and 405 (DAPI) channels.

We next investigated the results from the application of our method in more detail for the specific strains. From the genus Neisseria we selected a commensal strain, *N. lactamica*, and 12 pathogenic strains of *N. meningitidis. N. lactamica* is known to be present in the upper respiratory tract of humans (Dorey et al., 2019), and to confer natural immunity against meningititis in children. There are suggestions that the LOS could be implicated in this immunization (Braun et al., 2004; Gold et al., 1978; Kim et al., 1989; Maeland and Wedege, 1989; Sánchez et al., 2002). Sialylation of *N. lactamica* LOS has been reported, and the presence of the *lst* gene has been detected in several *N. lactamica* strains, including the one used in our study DSM 4691 Tsang et al. (2001), also detected the *lst* gene in all the *N. meningitidis* LOS prototype strains (L1-L12) used in this paper. We confirmed these results by PCR (Supplementary information, Table 3, Figure S9) using the primers designed by Tsang et al (Tsang et al., 2001).

Our data shows that *N. meningitidis* strains L1-L10 and L12, and serotypes B, C, W-135 and Y, and the commensal *N. lactamica*, are all capable of utilizing CMP-Neu5Az to incorporate sialic acid into their LOS, similar to *N. gonorrhoeae* (Figure 5A-C) (Jen et al., 2021). Strain L6 which lacks the terminal galactose residue could be sialylated, but to a lesser extent compared to other strains. This may indicate that the epitope found in the L6 strain LOS (4GlcNAcβ1–3Galβ1–4Glc) (Mubaiwa et al., 2017b) is not the most suitable substrate for its own sialyltransferase. Sialylation of strain L12 is also reduced compared to the others, however since the LOS of this strain is not fully elucidated yet we cannot hypothesize about the acceptor preference of the ST. Furthermore, we do not have information about the level of expression of the *lst* enzyme within the different strains. The lack of sialylation of the LOS of *N. meningitidis* L11 could also indicate the specificity of the enzyme for substrates containing Galβ1–4GlcNAcβ1–3Galβ1–4Glc, or to a lesser extent, 4GlcNAcβ1–3Galβ1–4Glc, but not for Glcβ1– 4Glc which is the epitope found in the L11 LOS (Mistreta et al., 2010). In the literature there is structural evidence of the presence of Neu5Ac in the LOS of *N. meningitidis* strains L1-L5 (Choudhury et al., 2008; Mandrell et al., 1991), strains L9-L12 belong to the serogroup A (Tsang et al., 2001), this group does not have Neu5Ac in the CPS, and this explains why sialic acid has not been found in the LOS of strains cultured without an external supply of Neu5Ac.

**Figure 5.**
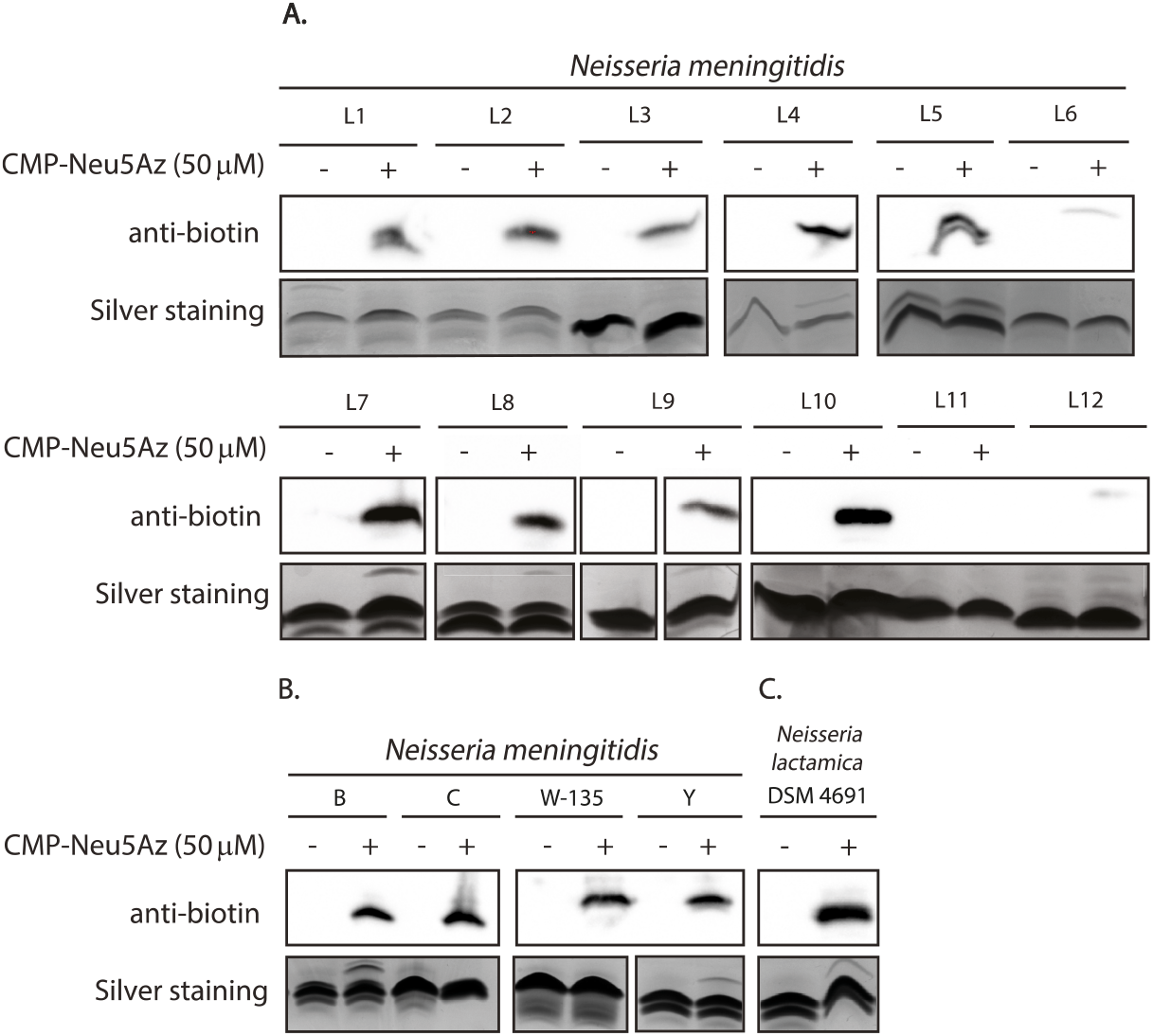
Scan for the sialyltransferase activity of *N. meningitidis* strains with CMP-Neu5Az. LOS was visualized on a silver-stained LOS/LPS gel, and sialylated LOS was detected in Western blot via SPAAC reaction with DBCO-PEG_4_-biotin.

From the genus Campylobacter, we observed labelling of different strains of *C. jejuni* GBS strains, *C. upsaliensis*, and *C. insulaenigreae*. Among these, *C. upsaliensis* and *C. insulaenigrae* are considered emerging zoonotic pathogens for humans (Chua et al., 2007; Fouts et al., 2005; Man, 2011; Richards et al., 2013a). Litle is known about the metabolism of Neu5Ac in these two bacteria, or the LOS structure that they express. Richards et al (2013) reported the genomic potential for sialic acid biosynthesis of strains of *C. upsaliensis* and *C. coli* (Richards et al., 2013b). To confirm the presence of these genes in our strain we designed primers targeting parts of the *CstII* gene based on the published *C. jejuni* sequence and found it is indeed present in *C. upsaliensis* and *C. insulaenigreae*. For the *C. coli* and *C. lari* strains in our lab, we did not detect the *CstII* genes, which is consistent with our LOS labelling results. Our labelling data showed that *C. upsaliensis* DMSZ 5365 and *C. insulaenigrae* DSM 17739 are indeed capable of transferring CMP-Neu5Ac onto their LOS, translating the genomic potential of these strains to functional evidence for a LOS active sialyltransferase and perhaps a role of sialylated LOS in its pathogenicity (Figure 6A).

**Figure 6.**
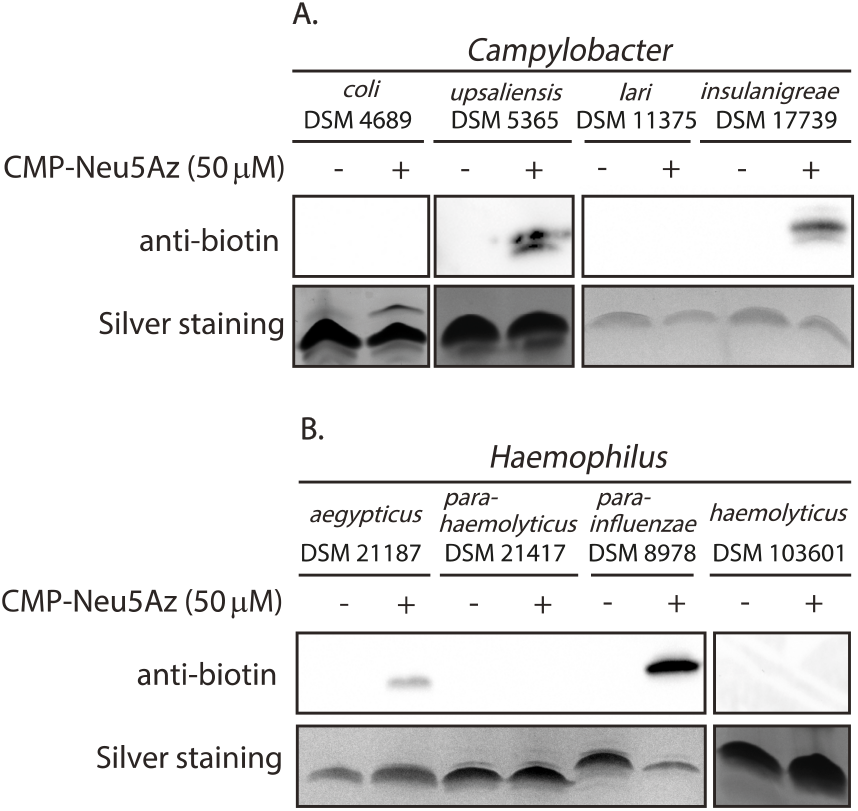
Scan for native ST activity of (A) Campylobacter strains and (B) Haemophilus strains with CMP-Neu5Az. Detected on a silver-stained LOS/LPS Western blot via SPAAC reaction with DBCO-PEG_4_-biotin.

For several Haemophilus species, it has been reported that their LOS can resemble human glycans (de Jong et al., 2022a). This form of glycan mimicry has also been observed for some strains of the pathogen *H. aegyptius* (Heise et al., 2018; Rubin, 1995; Swords et al., 2003). Sialylation of *H. aegyptius* LOS has however not been described yet, but we here show *H. aegyptius* DSM 21187 expresses active STs capable of introducing Neu5Ac onto its LOS (Figure 6B). This indicates it might perform a similar function in immune evasion as for non-typeable *H. influenzae* (Saha et al., 2021). For strains of commensal *H. parainfluenzae*, the LOS or LPS has been characterized and Neu5Ac was not reported to be present in these glycan structures (Pollard et al., 2008; Vitiazeva et al., 2011; Young and Hood, 2013). One report had scanned for the genetic potential to express an α2,3-sialyltransferase (*lic3A*), but could not identify this for any of the commensal bacteria (Young and Hood, 2013). Another report idenfied Neu5Ac as part of the O-antigen of strain 20 (Vitiazeva et al., 2011). Our results here indicate that *H. parainfluenzae* strain DSM 8978 expresses active STs that introduce Neu5Ac onto its LOS. We designed primers to assess the presence of genes for *Lic3A, LsgB* and *SiaA*, and consistent with previous findings, we were able to find it in *H. aegyptius* but not in *H. parainfluenzae*.

To test practical the applicability of our method to quickly type ST activity in clinical isolate collections, we screened 10 clinical non-typeable and 2 typeable isolates of *H. influenzae* using our protocol in combination with CMP-Neu5biotin. The LOS and sialylation ability of these non-typeable and typeable clinical isolates was unknown and we found that seven out of ten showed sialylation of their LOS, and a moderate signal was detected for both typeables strains (Figure 8). With this example, we demonstrate that this technique provides a rapid way to explore the sialyltransferase activity of multiple strains and to identify the sialylation of previously uncharacterized cell surface glycoconjugates of clinical isolates. Moreover, this NTHi sialyltransferase activity screen is also a rapid method for identifying functional sialylation of LOS, which provides a quick indication of whether the bacteria may use exogenous CMP-Neu5Ac *in vivo*.

We investigated the animal pathogen *P. multocida* serovar 3 strains Pm70 and P1059 that are known to display sialylated LOS/LPS (Harper et al., 2011), which is thought to shield the bacteria from the immune system (Tatum et al., 2009). *P. multocida* has a known sialyltransferase, *Pmst*, which also has sialidase activity (Yu et al., 2005). In our experiment, *P. multocida* DSM 5281 showed modest labelling of the LOS by nave sialyltransferases (Figure 7A). It is possible that the experimental conditions were not optimal for sialyltransferase activity, or that the sialidase activity of the enzyme also played a role. We tested another strain of the genus Pasteurella, *P. dagmatis*, an animal and human pathogen (Holst et al., 1992). This bacterium has a known sialyltransferase, *PdST*, which catalyzes the transfer of CMP-Neu5Ac onto the galactosyl residue of the LOS/LPS with an α2,3-linkage (Schmölzer et al., 2013). This enzyme hydrolyzes CMP-Neu5Ac in absence of an acceptor substrate (Schmölzer et al., 2014). In agreement with the suspected sialyltransferase activity, *P. dagmatis* showed labelling of its LOS/LPS.

**Figure 7.**
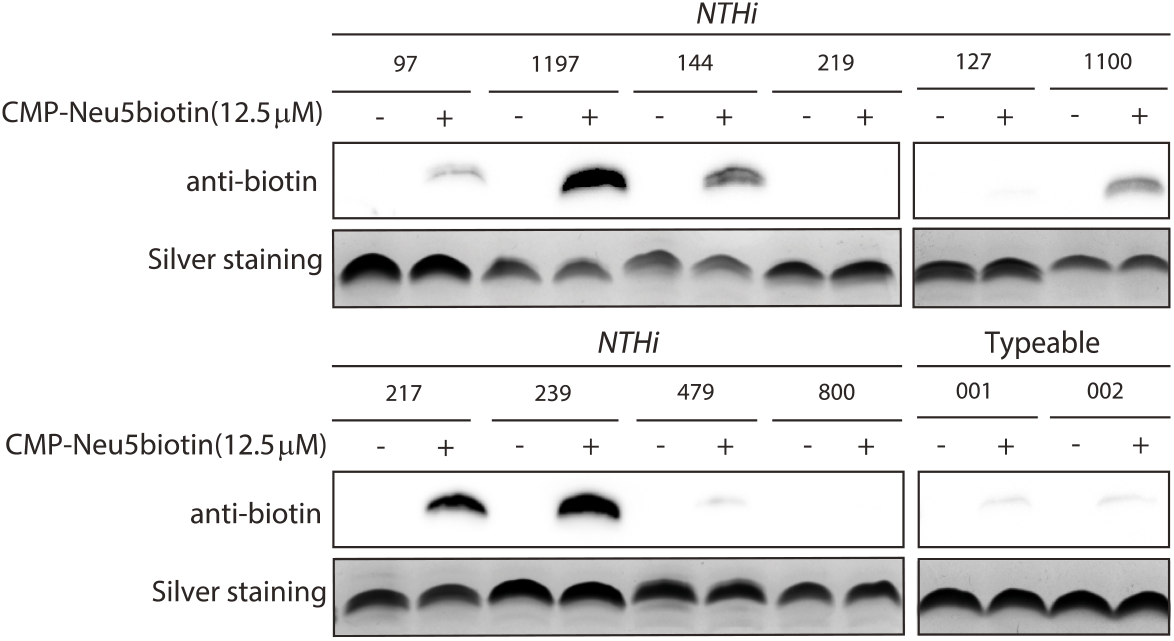
The native sialyltransferases of non-typeable and typeable *H. influenzae* strains show incorporation of CMP-Neu5Biotin to varying degrees, as observed after a strain-promoted click reaction with a DBCO-PEG_4_-biotin.

**Figure 8.**
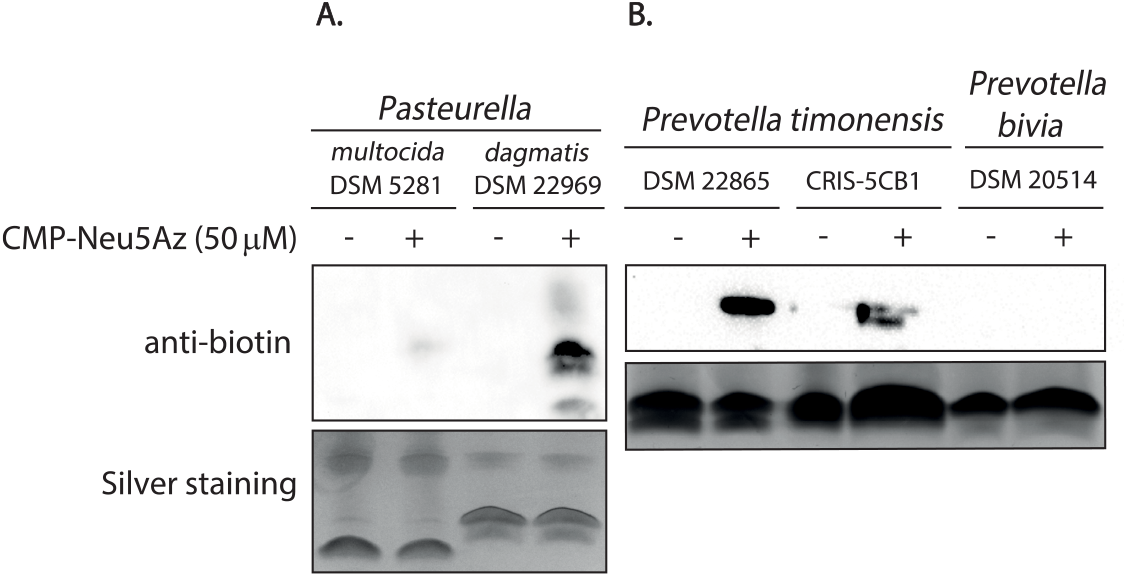
Scan for the sialyltransferase activity of (A) *Prevotella*, and (C) *Pasteurella* strains. CMP-Neu5Az was incorporated by native sialyltransferases followed by a strain-promoted click reaction with a DBCO-PEG_4_-biotin. The LOS/LPS was analyzed on Western blot and by silver stain.

Finally, we also studied two sialidase positive Prevotella strains *P. timonensis* 5C-1B and *P. bivia* both associated with bacterial vaginosis (Pelayo et al., 2024; Segui-Perez et al., 2024). Litle is known for these bacteria about their utilization of Neu5Ac, and their potential to display glycan mimicry (Srinivasan et al., 2015). By applying MOE, we found that strains of *P. timonensis* 5C-B1, TSW and ACES are capable of metabolizing and incorporating Neu5Az onto their LOS (Supplementary information, Figure S10). This suggests the presence of the enzymatic machinery capable of metabolizing Neu5Ac. Labelling experiments using CMP-Neu5Az also show that *P. timonensis* 5C-1B and *P. bivia* have active STs capable sialylating their LOS with exogenous CMP-Neu5Ac (Figure 7B). To the best of our knowledge, sialyltransferases from *P. timonensis* have not been described in the literature or have not been annotated in the genome yet. Taken together, these data suggest that Neu5Ac can be used by this bacterium to decorate its glycoconjugates, and we were able to assess this easily using our NST glycoengineering approach.

## Experimental section

### Material, methods and culturing conditions

Neu5Az, CMP-Neu5Az and CMP-Neu5Biotin, and CMP-Neu5AF_488_ were prepared as reported (de Jong et al., 2022b). HRP conjugated anti-biotin antibody (200-032-211) was purchased from Jackson ImmunoResearch Laboratories. Acetylene-PEG_4_-Biotin (CLK-TA105), DBCO-PEG_4_-Biotin (CLK-A105P4), AF_488_-alkyne (CLK-1277), and DBCO-AF_488_ (CLK-1278) were purchased from Jena Bioscience. Normal Human Serum (NHS) was a kind gi from UMC Utrecht. Multicolor low range protein ladder (26628) and ProLong Diamond Antifade Mountant (P36961) from, DAPI (4′,6-diamidino-2-phenylindole) (Merk), ECL Western Blotting Substrates (Bio-Rad Laboratories), Alcian Blue 8GX (Fluka, Sigma-Aldrich), DNA extraction kits (Macherey-Nagel GmbH & Co.), protein ladder and 1 Kb DNA ladder (Thermo Fisher Scientific), primers were ordered from Macrogen Europe. Agar blood sheep plates purchased from BioTrading (K004P090KP), chocolate plates (BioTrading, K018P090KP), chopped meat media and NYC media (DSMZ), trypticase soy yeast extract Oxoid™ (Thermo Fisher Scientific), heart infusion broth (BioTrading, K716F100GH), HEPES (Merk). The sialyltransferases CstII, Pmst1, Pmst3, Pd2,3-ST were expressed and purified in house (Utrecht University). All bacteria strains cultured as specified in the Supporting information. Further information on the bacterial strains used in this work can be found in Supporting information Table 1.

### SEEL of bacteria: (de Jong et al., 2022b)

One-step SEEL was performed on bacteria grown in liquid culture (5×10^8^ bacteria). The bacteria were washed with buffer and were incubated with SEEL label mix at 37°C for 2 h while rotating. A typical SEEL label mix (50μL) was prepared in medium (PBS/HEPES buffer) with sialyltransferase (1.05 uL of stock 1 mg/mL, or 2 mg/mL in case of Pmst3 and Pd2,3-ST), CMP-sialic acid derivative (50 μM), 0.34 uL BSA (2 mg/mL) and 0.34 μL alkaline phosphatase (1 U/ μL). After SEEL treatment, the bacteria were washed with buffer and prepared for application. Two-step SEEL was performed similar to one-step SEEL, followed by a click reaction. In case of CuAAC, 100 μL reaction volume contained 100 μM acetylene-PEG_4_-Biotin, 500 μM CuSO_4_ and 2.5 mM sodium L-ascorbate. In case of SPAAC 100 μL reaction volume contained 100 μM DBCO-PEG_4_-Biotin. For the fluorophores AF_488_-alkyne and DBCO-AF_488_ the concentration was 1 mM. If the bacteria were heat-inactivated, they were heated at 80°C for 15 min and treated with the described SEEL method.

### MOE with Neu5Az

Neu5Az were synthesized according to a published procedure (Han et al., 2005; Luchansky et al., 2004). MOE was performed with 6 × 10^8^ bacteria, and these were incubated with indicated concentrations of Neu5Az and Neu9Az. Bacteria were incubated for 6 h shaking at 160 rpm and for NTHi at 37°C, and for *C. jejuni* at 42°C under microaerophilic conditions. Aer incubation, the samples were washed 2 × 1 mL (buffer) and clicked via CuAAC or SPAAC, see SEEL of bacteria, washed again and further treated for (glyco)proteins or LOS.

### Labelling of bacterial LOS via nave sialyltransferases

Bacteria were grown as described previously. Bacteria (5×10^8^ CFU/mL) were centrifuged at 5000 rpm for 5 minutes and washed with HEPES (in case of *Neisseria*) or PBS. For two step labelling, bacterial pellet was resuspended in a solution of the sugar nucleotide (CMP-Neu5Az) in buffer at a final concentration of 50 μM, and incubated for 2 h at 37°C on rotating device. For negative controls bacteria were incubated with buffer. Bacteria, including negative controls, were washed 2 times with buffer and incubated 1 h with a solution of DBCO-PEG_4_-biotin 100 μM or DBCO-AF_488_, at room temperature. After this, bacteria were washed multiple times and resuspended in PBS. For one step labelling, we incubated the bacteria with a solution 50 μM of CMP-Neu5biotin or CMP-Neu5AF_488_ for two hours at room temperature. Bacteria were washed multiple times with PBS.

For LOS visualization samples were boiled for 5 min and then treated with proteinase K (10 μL, 20 mg/mL) overnight at 55°C.Laemmli buffer was added and loaded onto tris-tricine gels as described below. Western blot using an-Biotin HRP was employed to detect labelled LOS when CMP-Neu5Biotin was used or two steps labelling using CMP-Neu5Az followed by SPAAC with DBCO-PEG_4_-Biotin was performed. Loading controls were stained with the silver staining standard protocol described below.

In case of (glyco)proteins, the samples were lysed and analyzed with a 10% SDS-PAGE gel for which the gel was run for 45-60 min at 150 V. Western blots to detect Biotin tags and Page Blue to stain loading controls.

To check Sialylation of capsules (*N. meningitidis* B, C, W-135 and Y, and *C. jejuni* GB11 and GB19) 1×10^8^ bacteria was centrifuged at 13000 rpm for 5 minutes and resuspended in 60 µL of lysis buffer (0.32 mL 1M Tris/HCl pH6.8, 0.4 g SDS, 2.5 mg bromophenol blue, 2.0 mL glycerol, and 7.68 mL H_2_O). The supernatant was treated with 1.5 µL of a 20 mg/mL stock proteinase K, and heated at 50°C for 1h, followed by boiling at 100°C for 10 min. Samples were loaded on 10% SDS-Page gel, and ran at 125 V. Western blot was used to detect biotinylated CPS. For the loading controls, an adapted protocol of Lin was applied (Boleij et al., 2018). Briefly, after electrophoresis, gels were washed in a solution 25% ethanol and 10% acetic acid for 3.5 h. Gels were stained in 0.125% Alcian Blue in 25% ethanol and 10% acetic acid (pH 2.5) for one hour and followed by several washings with 25% ethanol and 10% acetic acid solution overnight, until bands appear.

### Western blong

For the western blotting, the gel was electrobloted onto a PVDF membrane. The membrane was blocked with 5% skimmed milk for 30 – 60 min, washed with 1% milk for 5 min, stained with anti-Biotin-HRP antibody (1:20000 in 1% milk), washed (1 % milk, followed by PBS, 5 min each), and treated with ECL Western substrate for signal detecon.

### LOS preparation and tris-tricine gel

Samples were loaded onto a 16% Tris-Tricine gel. The gel ran typically for 3 - 4 h at 20 mA and was then further analyzed by Western blotting and silver staining or in-gel fluorescence.

### Silver staining

Silver staining was performed on a 16% Tris-Tricine gel as described previously (Merril and Shifrin, 1982). Briefly, the gel was fixed (30 min, 40% ethanol, 5% acetic acid), oxidized (5 min, 0.7% sodium periodic acid, 40% ethanol, 5% acetic acid), washed (3 × 5 min in distilled water), stained (distilled water containing 19% 0.1 M NaOH, 1.3% >28% ammonium hydroxide, 3.3% 20% w/v silver nitrate), washed (3 × 5 min in distilled water), developed until bands appeared (distilled water containing 0.1% PFA 37% and 0.1% citric acid 100 mg/mL), rinsed with distilled water and stopped (7% acetic acid in distilled water).

### In-gel fluorescence

In-gel fluorescence was measured on Amersham imager 600 using the Green channel (520 nm, Cy3).

### Serum resistance assay

*N. gonorrhoeae* WT was incubated with and without 20 nmol/mL CMP-Neu5Ac or CMP-Neu5Az. After treatment, *N. gonorrhoeae* was washed and diluted to 10^6^ bacteria in HEPES buffer. Bacteria (1×10^4^) were treated with 10% NHS or HI-NHS for 1 h at 37°C. Samples were 25x diluted and 50 μL was plated out on chocolate columbia agar with vitox. Plates were grown at 37°C + 5% CO_2_ overnight, colony forming units were counted and data analysis was done with Prism software.

### Detection of genes by PCR

Bacteria were cultured as described previously. To extract chromosomal DNA, 1 mL of culture with an OD_600_ of 1 was centrifuged at 5000 rpm for 5 minutes. The DNA was extracted from the pellet using the instructions provided by a commercial kit (Macherey-Nagel GmbH & Co.). Each reaction consisted of 500 nM of primer FWD and REV (Table S3, supporting information), 200 nM of dNTP (Thermo Fisher Scientific), 5 U of DreamTaq DNA Polymerase (Thermo Fisher Scientific), and 1 μL of extracted DNA (15-25 μg/mL), in a total volume of 50 μL. PCR reactions were performed in a thermal cycler (MyCycler Bio-Rad) using a standard protocol. The cycling conditions consisted of an initial denaturation at 95°C for 3 minutes, followed by 30 cycles of denaturation at 95°C for 30 seconds, annealing at the appropriate temperature for 30 seconds, and extension at 72°C for 45 seconds, with a final extension period of 5 minutes. The amplification products were visualized by electrophoresis on a 1% agarose gel and Midori green staining, and then detected using a ChemiDoc MP system (Bio-rad). PCR products were sent to sequence without purification.

### Fluorescence confocal microscopy

Bacteria (circa 3×10^8^ cells) were centrifuged, resuspended in HEPES + 1 % BSA and DAPI (4′,6-diamidino-2-phenylindole) (1:50) was added. The samples were incubated for 25 min in the dark before washing (milliQ + 0.1% Tween) and then carefully resuspended in ProLong Diamond Antifade Mountant (P36961). 10 μL of sample was taken, put on a poly-L-lysine coated coverslip and mounted on a glass slide. Slides were stored at RT overnight to allow the samples to harden. Images were collected on Olympus/Evident SpinSR10 confocal microscope in combination with SORA software. Image analysis was performed using OliVia/ImageJ software.

## Conclusion

Using an expansive and diverse set of bacteria, with many relevant bacteria to human health and disease, we here demonstrate that glycoengineering with native sialyltransferases offers a straightforward, quick, and non-toxic approach for adding various modifications to the lipooligosaccharides (LOS) of bacteria. This labelling protocol also enables the easy and quick assessment of the presence of active bacterial sialyltransferases and their potential to sialylate its LOS with exogeneous CMP-Neu5Ac, regardless of their ability to produce and metabolize Neu5Ac. The localization of the sialyltransferases is a question we aim to address in future studies, however, the rapid labeling time we observed using CMP-Neu5Az suggests the presence of sialyltransferases near or on the outer surface of the bacteria, as we have shown for *N. gonorrhoeae*. Using our method, we annotate and validate for the first time for a wide array of bacterial pathogens that they exhibit LOS sialyltransferase activity, which opens avenues for more in-depth exploration of the glycosylation processes in bacterial LOS. This method can also aid the discovery of new sialyltransferases with relaxed donor specificity and significant potential for their use in chemoenzymatic synthesis. Our method can be used to introduce modified sialic acid onto bacterial LPS/LOS, allowing precise labelling, visualization and tracking of Neu5Ac on bacterial surfaces via click chemistry. We show that methods such as MOE, SEEL, and labelling through native sialyltransferases serve as complementary glycoengineering techniques, with the choice among them depending on the specific objectives. This approach provides an additional tool to the existing strategies available for the study of the role of Neu5Ac in bacteria.

## Author contribution

E.I.A.M.: Writing – Original Draft, Review and Editing, Conceptualization, Methodology, Resources, Investigation, Validation and Formal Analysis. H.J.: Writing, Review and Editing, Conceptualization, Methodology, Resources, Investigation, Validation and Formal Analysis. J.E.H.: Investigation. J.Y.O.: Resources. M.M.S.M.W.: Conceptualization, Review and Editing, Formal Analysis, Supervision. T.W.: Conceptualization, Writing, Review and Editing, Formal Analysis, Supervision, Funding acquisition and Project administration.

## Funding

This work was supported by the Dutch Research Council (NWO) via a VIDI grant (723.014.005 to TW) and BBOL grant (737.016.013). Funding from the European Union’s Horizon 2020 Marie Skłodowska-Curie Actions for the Innovative Training Network “Sweet Crosstalk” under the grant agreement No 814102.

## Acknowledgements

We thank Prof. dr. Geert-Jan Boons and dr. Gerlof P. Bosman for providing the recombinant sialyltransferases used in the SEEL experiments. Prof. dr. Nina van Sorge and the Netherlands Reference Laboratory for Bacterial Meningitis (NRLBM) for generously providing the *Neisseria meningitidis* strains L1-L12. Dr. Karin Strijbis and Celia Segui-Perez for providing the Prevotella strains described in this study. Nontypeable *Haemophilus influenzae* R2886 and the *siaP* mutant were a kind gift from dr. Jeroen Langereis, Radboudumc. Other nontypeable *Haemophilus influenzae* and typeable strains were a kind gift from Clinical Infectiology, Utrecht University. Normal Human Serum (NHS) was a kind gift from prof. dr. Suzan Rooijakkers from UMC Utrecht. Dr. Astrid Heikema from Erasmus MC Rotterdam for providing *C. jejuni* GB strains and their *cstII* mutants. Prof. dr. Jos van Putten for the *Neisseria gonorrhoeae* and *Neisseria meningitidis* serogroup B, C, W-135 and Y strains. Dr. Maria J. Moure for the Sia-AF488-CMP used in this study.

## Conflict of Interest

The authors declare no conflict of interest.

## Notes

### Competing Interest Statement

The authors have declared no competing interest.

