## Supporting Information for "Glycoengineering with neuraminic acid analogs to label lipooligosaccharides and detect native sialyltransferase activity in Gram-negative bacteria"

**A glycoengineering strategy for labelling lipooligosaccharides and detecting native sialyltransferase activity in live Gram-negative bacteria with neuraminic acid analogs**

Erianna I. Alvarado Melendez [a,#], Hanna de Jong [a,#], Jet E. M. Hartman [a], Jun Yang Ong [a], Marc M. S. M. Wösten [b*] and Tom Wennekes [a*]

**Supporting information:**

**Table S1.** Bacterial strains used in this work.

| Species | Strain | Labelling NSTs | PCR primers | PCR product detected |
|---|---|---|---|---|
| *Campylobacter coli* | DSM 4689 | + | | |
| *Campylobacter insulaenigrae* | DSM 17739 | + | CstII cust | + |
| *Campylobacter jejuni* | GB 2 | + | CstII | + |
| *Campylobacter jejuni* | GB 2 ΔCstII | | CstII | |
| *Campylobacter jejuni* | GB 11 | + | CstII | + |
| *Campylobacter jejuni* | GB 11 ΔCstII | | CstII | |
| *Campylobacter jejuni* | GB 19 | + | CstII | + |
| *Campylobacter jejuni* | GB 19 ΔCstII | | CstII | |
| *Campylobacter lari* | DSM 11375 | - | | |
| *Campylobacter upsaliensis* | DSM 5365 | - | CstII cust | + |
| *Haemophilus aegyptius* | DSM 21187 | + | Lic3A, LsgB, SiaA | + |
| *Haemophilus haemolyticus* | DSM 103601 | - | LsgB | + |
| *Haemophilus parahaemolyticus* | DSM 21417 | - | | |
| *Haemophilus parainfluenzae* | DSM 8978 | + | | |
| *Helicobacter pullorum* | DSM 23160 | - | | |
| *Neisseria gonorrhoeae* | F62 | + | Lst3, Lst4 | |
| *Neisseria gonorrhoeae* | F62 Δlst | | | |
| *Neisseria lactamica* | DSM 4691 | + | Lst3, Lst4 | |
| *Neisseria meningitidis L1* | 126E | + | Lst3, Lst4 | |
| *Neisseria meningitidis L2* | 35E | + | Lst3, Lst4 | + |
| *Neisseria meningitidis L3* | 6275 | + | Lst3, Lst4 | + |
| *Neisseria meningitidis L4* | 891 | + | Lst3, Lst4 | + |
| *Neisseria meningitidis L5* | M981 | + | Lst3, Lst4 | + |
| *Neisseria meningitidis L6* | M992 | + | Lst3, Lst4 | + |
| *Neisseria meningitidis L7* | 6155 | + | Lst3, Lst4 | + |
| *Neisseria meningitidis L8* | M978 | + | Lst3, Lst4 | + |
| *Neisseria meningitidis L9* | 120M | + | Lst3, Lst4 | + |
| *Neisseria meningitidis L10* | 7880 | + | Lst3, Lst4 | + |
| *Neisseria meningitidis L11* | 7889 | - | Lst3, Lst4 | + |
| *Neisseria meningitidis L12* | 7897 | + | Lst3, Lst4 | + |
| *Neisseria meningitidis serogroup B* | | + | Lst3, Lst4 | + |
| *Neisseria meningitidis serogroup C* | | + | Lst3, Lst4 | + |
| *Neisseria meningitidis serogroup W-135* | | + | Lst3, Lst4 | + |
| *Neisseria meningitidis serogroup Y* | | + | Lst3, Lst4 | + |
| *Nontypeable Haemophilus influenzae* | R2886 | + | | |
| *Nontypeable Haemophilus influenzae* | R2886 ΔSiaP | | | |

| | | | | |
|---|---|---|---|---|
| *Nontypeable Haemophilus influenzae* | 97 | + | | |
| *Nontypeable Haemophilus influenzae* | 1197 | + | | |
| *Nontypeable Haemophilus influenzae* | 144 | + | | |
| *Nontypeable Haemophilus influenzae* | 219 | - | | |
| *Nontypeable Haemophilus influenzae* | 127 | + | | |
| *Nontypeable Haemophilus influenzae* | 1100 | + | | |
| *Nontypeable Haemophilus influenzae* | 217 | + | | |
| *Nontypeable Haemophilus influenzae* | 239 | + | | |
| *Nontypeable Haemophilus influenzae* | 479 | + | | |
| *Nontypeable Haemophilus influenzae* | 800 | - | | |
| *Pasteurella dagmatis* | DSM 22969 | + | Pmst1 | + |
| *Pasteurella multocida* | DSM 5281 | + | Pmst1 | + |
| *Photobacterium leiognathid* | DSM 21260 | - | | |
| *Prevotella bivia* | DSM 20514 | - | | |
| *Prevotella timonensis* | CRIS 5C-B1 | + | | |
| *Prevotella timonensis* | DSM 22865 | + | | |
| *Typeable Haemophilus influenzae* | 001 | + | | |
| *Typeable Haemophilus influenzae* | 002 | + | | |

**Table S2.** Blast search of bacterial sialyltransferases.

| Organism/ST | Locus | Amino acid length | Amino acid similarity |
|---|---|---|---|
| ***Neisseria gonorrhoeae/F62*** | AAY41922 | 378 | 100 % (378/378) |
| ***Neisseria meningitidis*** | WP_096137355 | 378 | 374/378(99%) |
| ***Neisseria lactamica*** | WP_118846604 | 378 | 364/378(96%) |
| *Neisseria polysaccharea* | WP_304673934 | 378 | 362/378(96%) |
| *Neisseria bergeri* | WP_304671870 | 378 | 360/378(95%) |
| *Neisseria viridiae* | WP_301911444 | 371 | 359/371(97%) |
| *Neisseria blantyrii* | WP_304671466 | 371 | 357/371(96%) |

| Organism/ST | Locus | Amino acid length | Amino acid similarity |
|---|---|---|---|
| ***Haemophilus influenzae R2866/Lic3A*** | ADO80211 | 326 | 100 % (326/326) |
| ***Haemophilus aegyptius*** | OBX78592 | 326 | 303/330(92%) |
| ***Haemophilus haemolyticus*** | WP_118795855 | 318 | 297/325(91%) |
| *Streptococcus pneumoniae* | CVP26011 | 275 | 255/323(79%) |
| *Haemophilus felis* | NBI40444 | 344 | 167/298(56%) |

| Organism/ST | Locus | Amino acid length | Amino acid similarity |
|---|---|---|---|
| ***Haemophilus influenzae/SiaA*** | ADO81445.1 | 304 | 100 % (304/304) |
| ***Haemophilus haemolyticus*** | WP_118802758.1 | 304 | 290/304(95%) |

| | | | |
|---|---|---|---|
| ***Haemophilus parainfluenzae*** | WP_255308331.1 | 304 | 231/303(76%) |
| ***Haemophilus parahaemolyticus*** | WP_258789035.1 | 304 | 227/303(75%) |
| ***Haemophilus aegyptius*** | WP_257003955.1 | 218 | 215/218(99%) |
| *Pasteurellaceae* | WP_210432889.1 | 304 | 201/303(66%) |
| *Rodentibacter heylii* | WP_059367401.1 | 304 | 201/303(66%) |

| Organism/ST | Locus | Amino acid length | Amino acid similarity |
|---|---|---|---|
| ***Haemophilus influenzae/LsgB*** | WP_179224786.1 | 304 | 100 % (304/304) |
| ***Haemophilus haemolyticus*** | WP_118793947.1 | 304 | 289/304(95%) |
| *Streptococcus pneumoniae* | CVP80484.1 | 304 | 287/304(94%) |
| ***Haemophilus aegyptius*** | WP_065251432.1 | 304 | 282/304(93%) |

| Organism/ST | Locus | Amino acid length | Amino acid similarity |
|---|---|---|---|
| ***Campylobacter jejuni/CstI*** | AWB40613 | 294 | 100 % (294/294) |
| ***Campylobacter lari*** | WP_214125788.1 | 294 | 291/294(99%) |
| ***Campylobacter coli*** | ECK0302889.1 | 236 | 236/236(100%) |
| *Campylobacter subantarcticus* | WP_039664428.1 | 300 | 183/304(60%) |
| ***Campylobacter insulaenigrae*** | WP_257908147.1 | 294 | 168/292(58%) |
| *Campylobacter sp.* | WP_291953368.1 | 291 | 163/288(57%) |
| *Campylobacter volucris* | TXE89740.1 | 247 | 146/247(59%) |

| Organism/ST | Locus | Amino acid length | Amino acid similarity |
|---|---|---|---|
| ***Campylobacter jejuni/CstII*** | ABN43105 | 291 | 100 % (291/291) |
| ***Campylobacter coli*** | EHJ4415117.1 | 291 | 260/288(90%) |
| *Escherichia coli* | GJJ32339.1 | 291 | 258/288(90%) |
| *Campylobacter bilis* | WP_066776435.1 | 296 | 228/289(79%) |
| *Campylobacter hepaticus* | MDC5558492.1 | 245 | 210/290(72%) |
| *Acinetobacter baumannii* | WP_210432889.1 | 304 | 208/245(85%) |
| *Campylobacter canadensis* | WP_172231115.1 | 301 | 167/295(57%) |
| ***Campylobacter lari*** | WP_257937707.1 | 296 | 164/293(56%) |
| ***Campylobacter insulaenigrae*** | WP_257938339.1 | 294 | 160/292(55%) |
| *Campylobacter subantarcticus* | WP_039664428.1 | 300 | 166/290(57%) |

| Organism/ST | Locus | Amino acid length | Amino acid similarity |
|---|---|---|---|
| ***Pasteurella multocida/Pmst1*** | AAK02272 | 412 | 100 % (412/412) |
| ***Pasteurella dagmatis*** | AWW59956.1 | 412 | 411/412(99%) |
| *Pasteurella oralis* | WP_101774701.1 | 412 | 296/412(72%) |
| *Escherichia coli* | WP_213061541.1 | 240 | 236/240(98%) |
| *Mergibacter septicus* | WP_265474447.1 | 425 | 204/414(49%) |
| *Shewanella halifaxensis* | WP_108944675.1 | 514 | 155/395(39%) |
| *Photobacterium leiognathi* | BAI49484.1 | 511 | 156/395(39%) |

**Table S3.** Primers sequences used in this study.

| **Primers** | **A (FWD)** | **B (REV)** |
|---|---|---|
| **CstII** | 5’-GAG ACC GAA CTA ATC ATG TG-3’ | 5’-TTT GTA TAG GGC TAT GG-3’ |
| **SiaA** | 5’-GAA GAT GGC ACT GAG AAC TAT C-3’ | 5’-GGG TAT ACA TAT AC GCC GTT GAG-3’ |
| **Ng Lst** | 5’-GGT GGC AGA AAG GAT TAT GG-3’ | 5’-CCC GTA TCA GGT TGT TTG TG-3’ |
| **LsgB** | 5’-CGT CAT GTT TGG TGT ATG G -3’ | 5’-CGG TAA TCT TCT GCT GGA TGA G-3’ |
| **Lic3A** | 5’-GGG AAT CCT TAC GCA TTT CAT C-3’ | 5’-AGA GGA CTT TCT GGC GAA ATA C-3’ |

| | | |
|---|---|---|
| **Pmst1** | 5'-GCT CGT TAT GTC TGG CAA TC-3' | 5'-GCA ACA CCA CCC ACT TTA TC-3' |
| **Cst cust** | 5'-AAT CAT TTT ATT TTG AAG ATA AAT A-3' | 5'-CGG CAA TTG CAC ACA TAT A-3' |
| **Lst3*** | 5'-CGT TAA ATT CGC CCT ATG TCA-3' | 5'-GGG CAT ACA CCG GCT TGA GCC-3' |
| **Lst5*** | 5'-CGT TAA ATT CGC CCT ATG TCA-3' | 5'-ATC GGG ATG CCG GAT TGG GTA-3' |

(*) From reference: R.S.W. Tsang, D.K.S. Law, C.M. Tsai, L.K. Ng, Detection of the lst gene in different serogroups and LOS immunotypes of Neisseria meningitidis, *FEMS Microbiol. Lett.* 199 (2001) 203–206. https://doi.org/10.1016/S0378-1097(01)00189-6.

### Supplementary information on bacterial strains and growing conditions:

*Campylobacter jejuni* GB11, and GB19, wildtype and mutants were a kind gift from Astrid Heikema, ErasmusMC Rotterdam. *Neisseria meningitidis* L1-12 were a kind gift from Nina van Sorge, Amsterdam UMC and the Netherlands Reference Laboratory for Bacterial Meningitis (NRLBM). *Neisseria gonorrhoeae* and *Neisseria meningitidis* serogroup B, C, W-135 and Y were a kind gift from Jos van Putten, Utrecht University. Nontypeable *Haemophilus influenzae* R2886 and the *SiaP* mutant were a kind gift from Jeroen Langereis, Radboudumc. Other nontypeable *Haemophilus influenzae* and typeable strains were a kind gift from Clinical Infectiology, Utrecht University. Prevotella strains were a kind gift from Karin Strijbis, Infection and Immunity Utrecht University. Other strains were purchased from DSMZ-German Collection of Microorganisms and Cell Cultures GmbH. Bacteria were cultured and grown under the following conditions:

***Campylobacter* species:**

*Campylobacter coli* and *Campylobacter lari* were grown on blood agar sheep plates (BioTrading, K004P090KP) at 37°C under microaerophilic conditions (80% $N_2$, 10% $CO_2$, 5% $O_2$, 5% $H_2$). *Campylobacter insulaenigrae* and *Campylobacter upsaliensis* were grown under the same conditions for 2 days. *Campylobacter jejuni* strains were grown under microaerophilic conditions at 42°C with a second passage for the mutant strains to allow sufficient growth. *Campylobacter coli*, *insulaenigrae*, *lari* and *upsaliensis* were grown at 37°C and *Campylobacter jejuni* at 42°C under microaerophilic conditions in Heart Infusion broth (BioTrading, K716F100GH).

***Haemophilus* species:**

*Haemophilus aegyptius*, *Haemophilus haemolyticus*, *Haemophilus parahaemolyticus*, *Haemophilus parainfluenzae* and typeable *Haemophilus influenzae* were grown on chocolate agar plates with Vitox (Thermo Scientific, PO5090A 0A) at 37°C with 5% $CO_2$ for 24 h. Nontypeable *Haemophilus influenzae* was grown aerobically. *Haemophilus* strains were cultured in Mueller Hinton or Brain Heart infusion supplemented with *Haemophilus* test supplement (Oxoid, SR0158E), except for *Haemophilus haemolyticus* which was cultured in tryptone soy broth. All strains were grown at 37°C for 24h.

***Helicobacter pullorum*** was grown on blood agar sheep plates (BioTrading, K004P090KP) at 37°C under microaerophilic conditions for 24 h (80% $N_2$, 10% $CO_2$, 5% $O_2$, 5% $H_2$) and in Heart Infusion broth (BioTrading, K716F100GH).

***Neisseria* species:**

*Neisseria gonorrhoeae, lactamica, and meningitidis* were cultured on chocolate columbia agar (BioTrading, K018P090KP) at 37°C with 5% $CO_2$ and in HEPES medium at 37°C for 24h.

***Pasteurella* species:**

*Pasteurella multocida* and *Pasteurella dagmatis* were grown on blood agar sheep plates (BioTrading, K004P090KP) at 37°C for 24 h, and in tryptone soy broth at the same temperature and incubation time.

***Prevotella* species:**

Cultures of *Prevotella bivia* and *Prevotella timonensis* were grown in New York City (NYC) medium and Cooked Meat Medium (CMM) respectively, supplemented with vitamin K1 (final concentration 1 mg/L) and hemin (final concentration 5 mg/L). Bacteria were cultured overnight at 37°C under anaerobic conditions in a Coy Vinyl Anaerobic Chamber. The $OD_{600}$ was measured on Biowave CO8000 Cell Density Meter.

***Photobacterium leiogonathid*** was cultured in marine broth at 30°C for 24 h.

**Supplementary Figure 1.** Protein gels of to NTHi WT and the Δ*SiaP* mutant after MOE treatment.

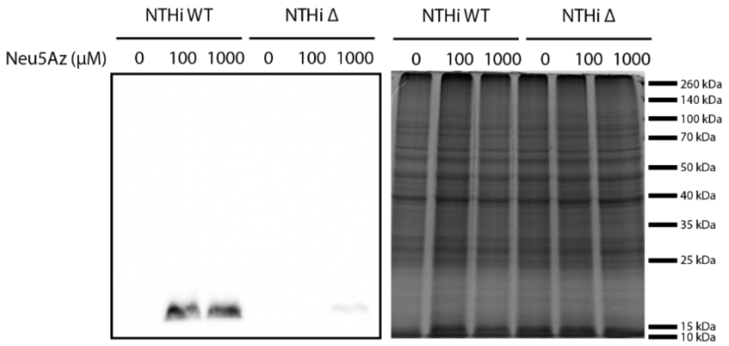


Anti-biotin Page Blue

**Supplementary Figure 2**. NTHi growth curves in the presence of Neu5Az (A) NTHi WT and (B) NTHi Δ*SiaP*.

Bacteria were treated according to conditions specified in text. Bacteria were diluted to $OD_{600}$ = 0.05 and the growth was monitored with Synergy HTX multi-mode meter in a hypoxic glove box for 24 h while shaking continuously. Data was exported and analyzed with excel or prism.

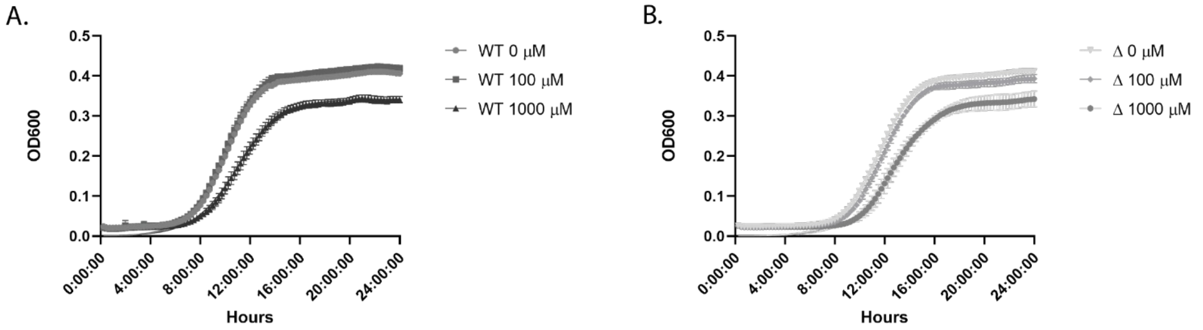


**Supplementary Figure 3**. SEEL applied to **NTHi using different exogenous sialyltransferases.**

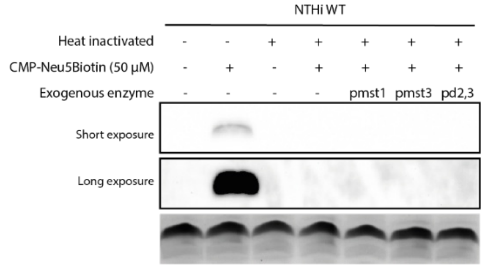


**Supplementary Figure 4.** SEEL applied to *C. jejuni* GB19.

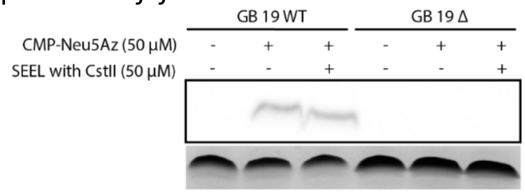

**Supplementary Figure 5.** SEEL applied to *C. jejuni* GB19, *CstII* and *CstI.*

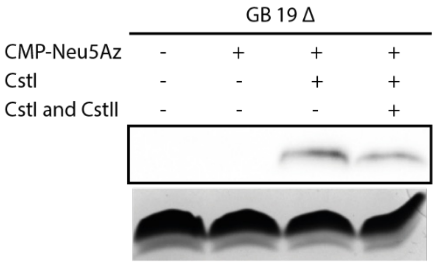


**Supplementary Figure 6.** Native sialyltransferase labelling on *N. gonorrhoeae* WT and Δ*lst* mutant followed by click with alkyne-$PEG_4$-biotin (CuAAC) vs. DBCO-biotin (SPAAC).

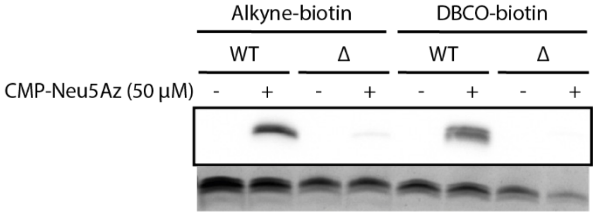

**Supplementary Figure 7.** Full length protein gels.

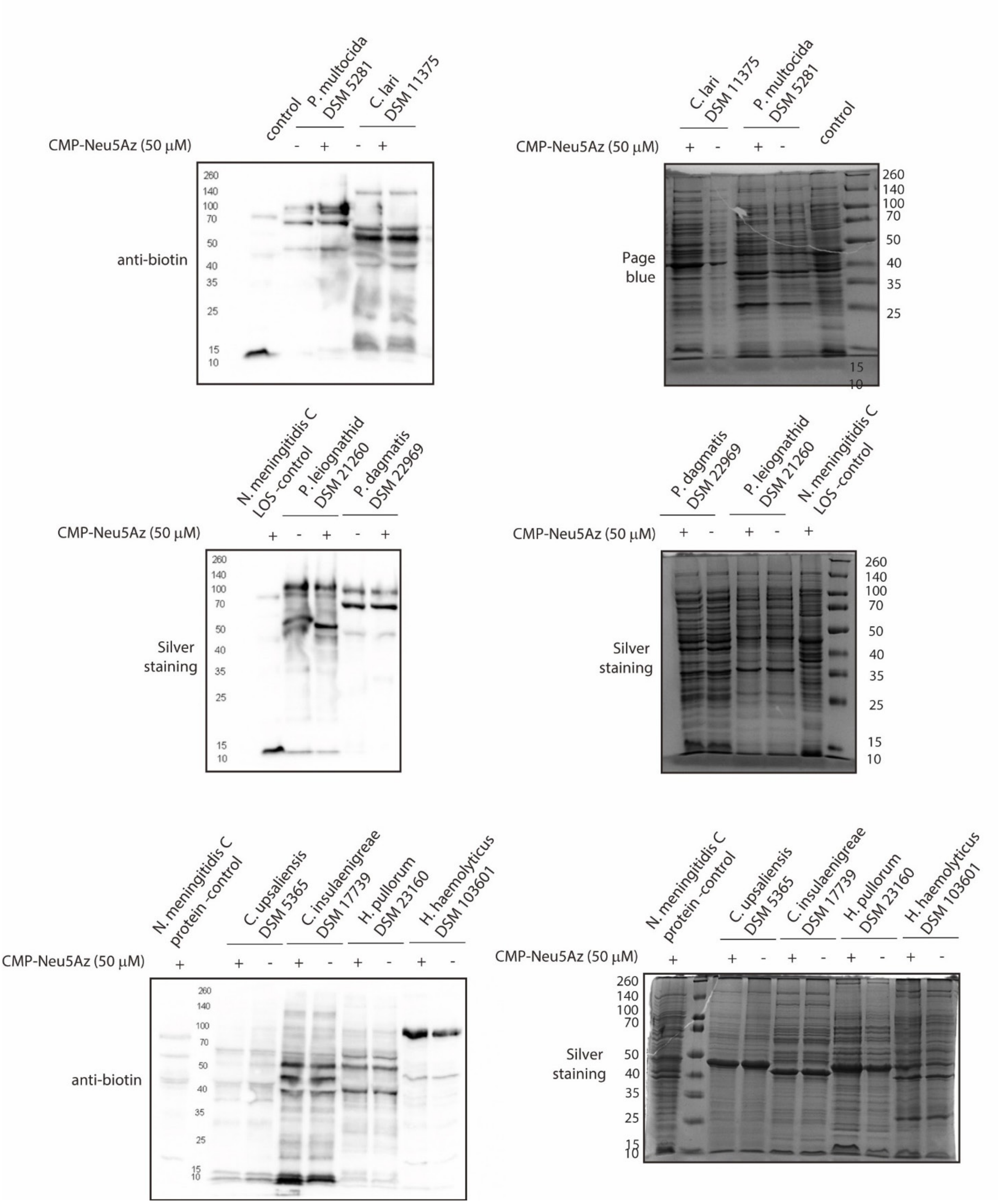

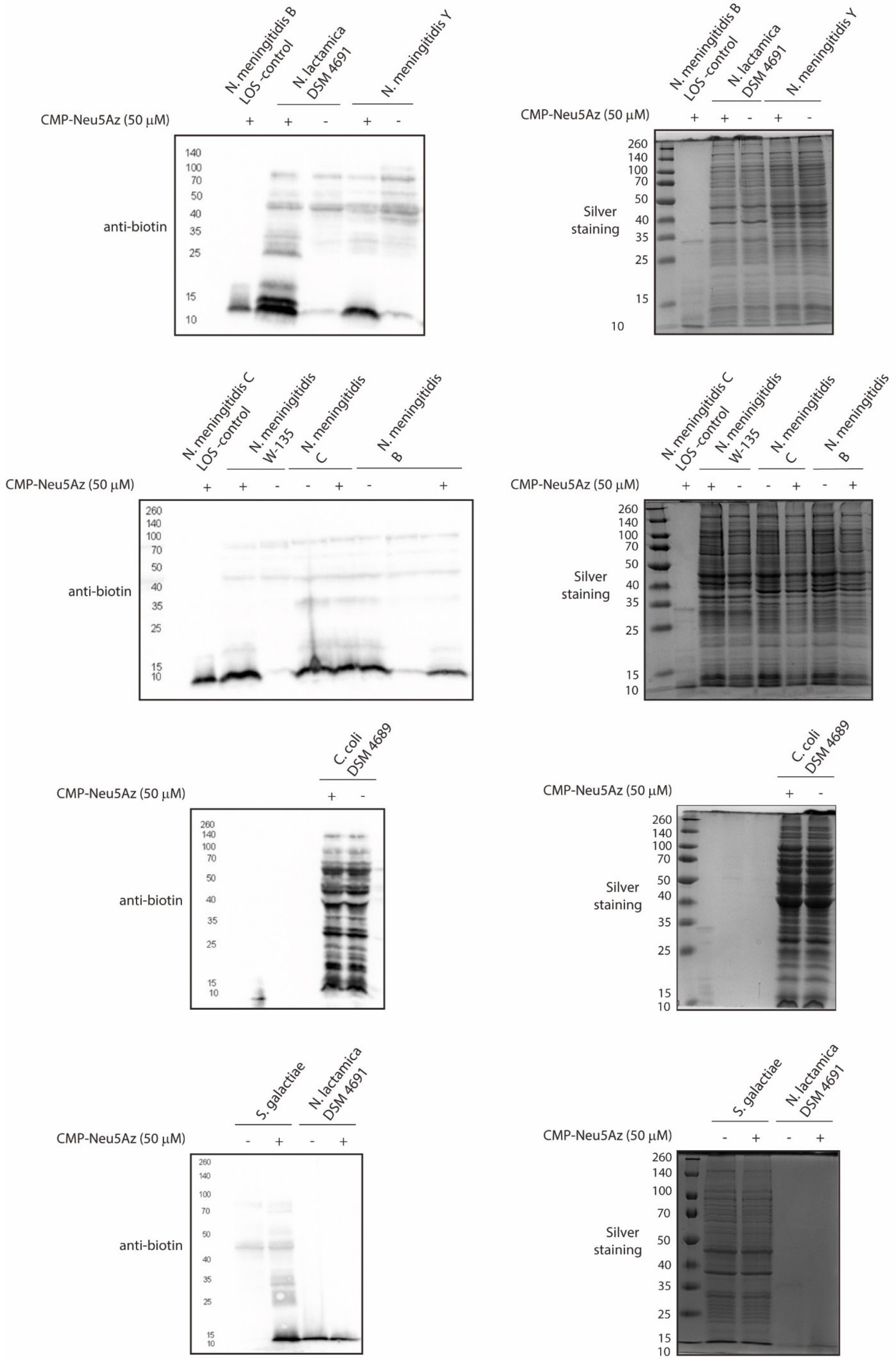
N. meningitidis B LOS-control
N. lactamica DSM 4691
N. meningitidis Y
CMP-Neu5Az (50 μM)
anti-biotin
Silver staining
N. meningitidis C LOS-control
N. meningitidis W-135
N. meningitidis C
N. meningitidis B
C. coli DSM 4689
S. galactiae
260
140
100
70
50
40
35
25
15
10

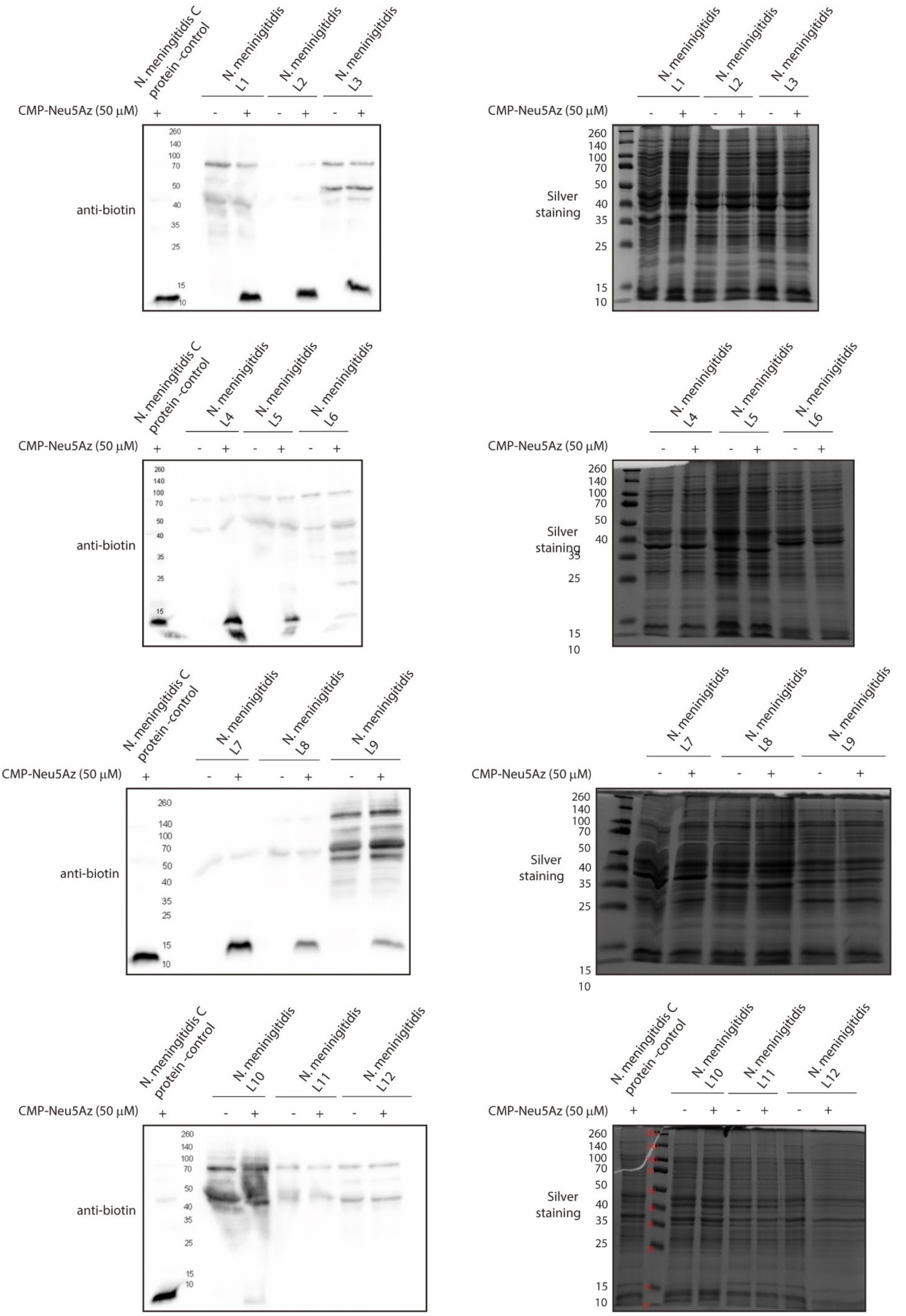
N. meningitidis C protein -control
N. meningitidis L1
N. meningitidis L2
N. meningitidis L3
CMP-Neu5Az (50 μM)
+ - + - + - +
anti-biotin
260
140
100
70
50
40
35
25
15
10
Silver staining
N. meningitidis L4
N. meningitidis L5
N. meningitidis L6
N. meningitidis L7
N. meningitidis L8
N. meningitidis L9
N. meningitidis L10
N. meningitidis L11
N. meningitidis L12

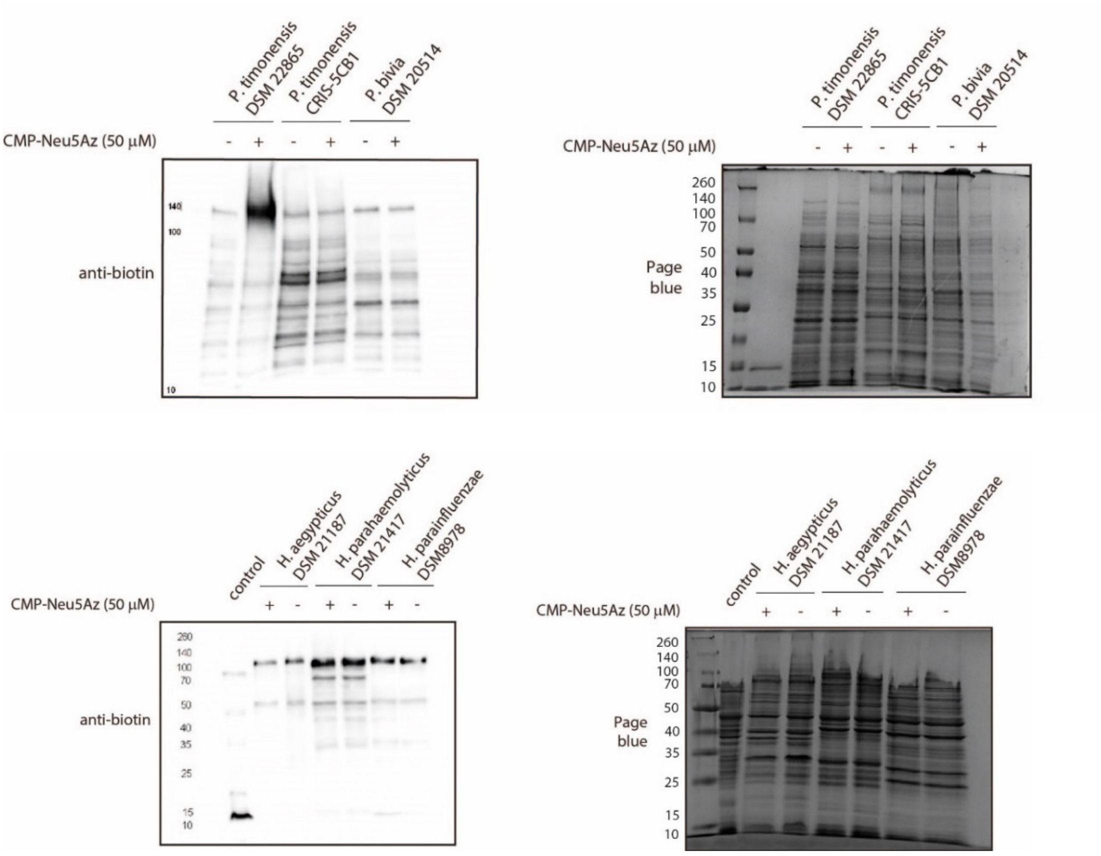


**Supplementary Figure 8.** CPS gels.

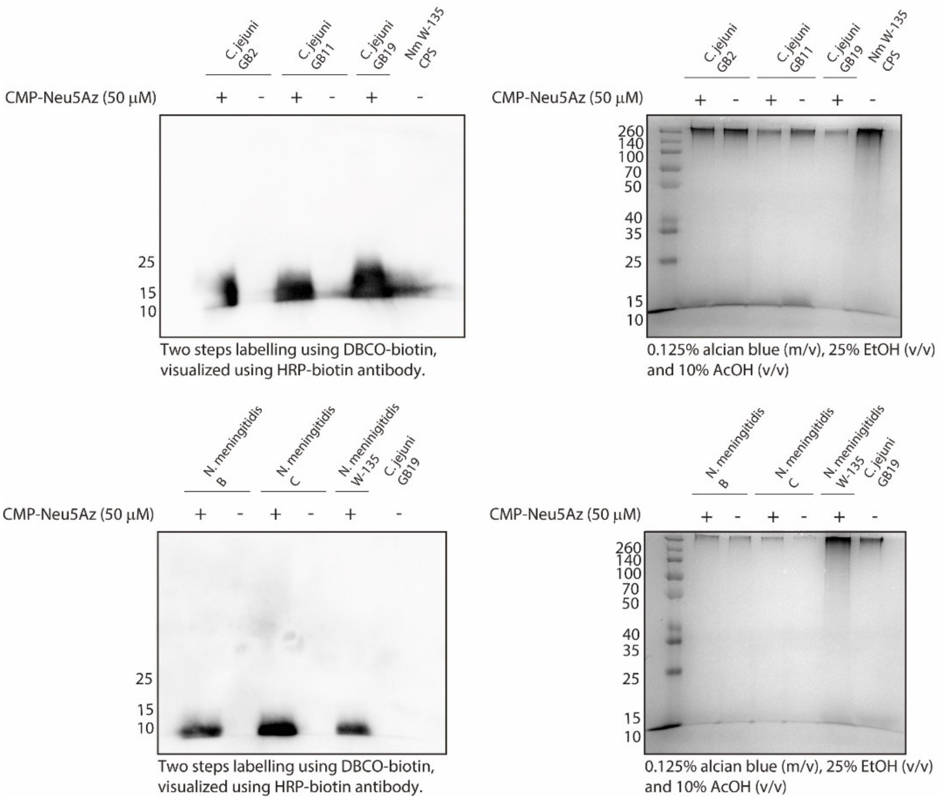

**Supplementary Figure 9.** PCR agarose gel for detection of PCR products.

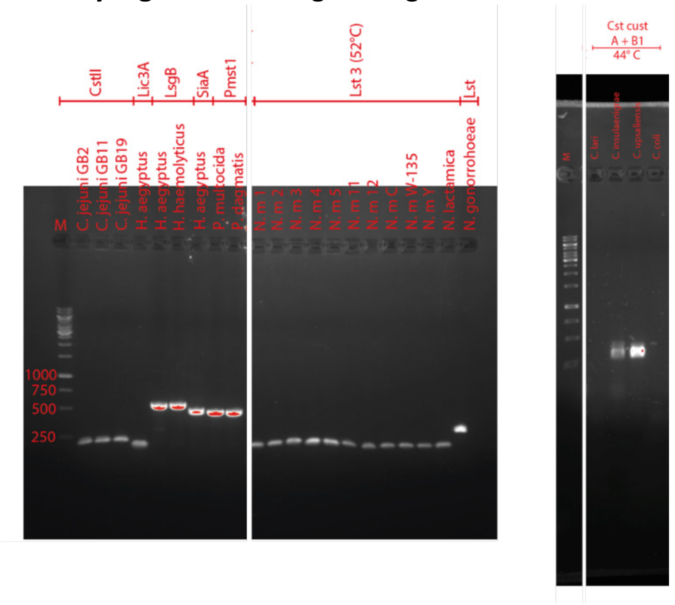


**Supplementary Figure 10.** MOE applied to *N. lactamica*, *N. meningitidis L12*, *P. dagmatis*, *P. bivia*, and various strains of *P. timonensis.*

**LOS gels:**

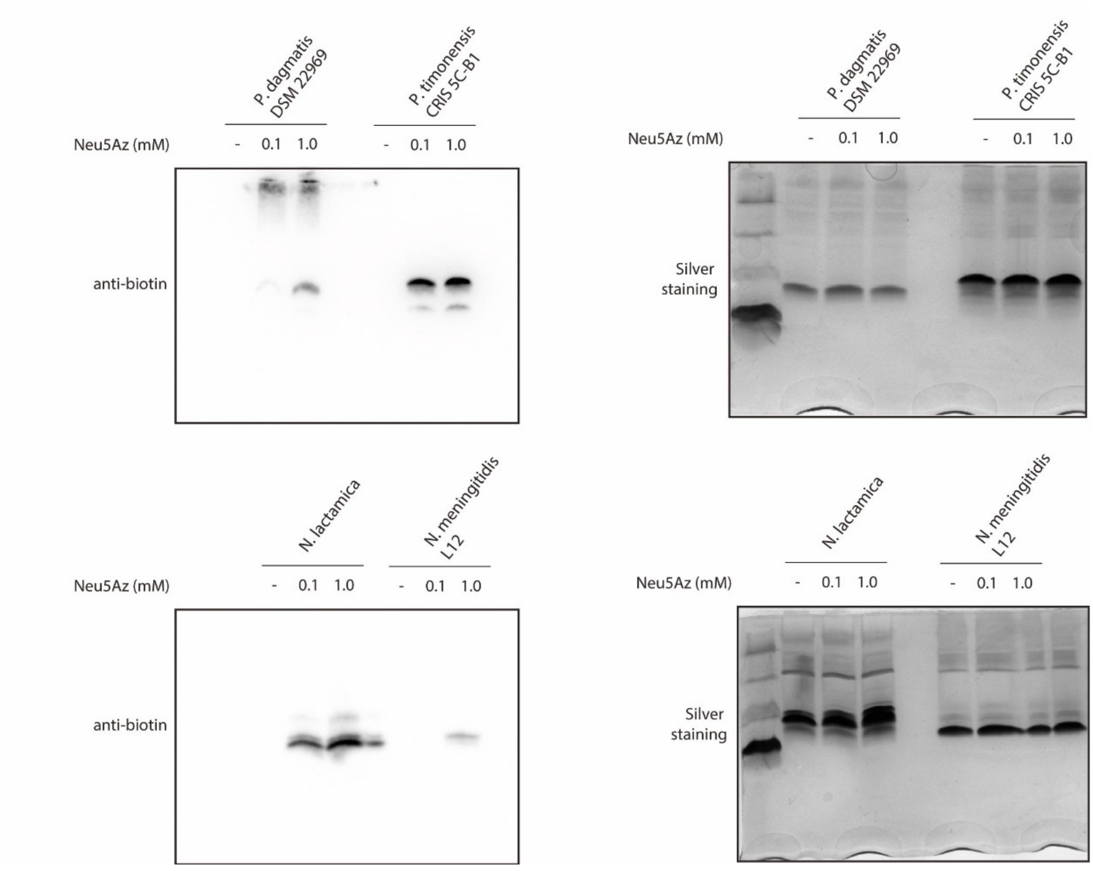

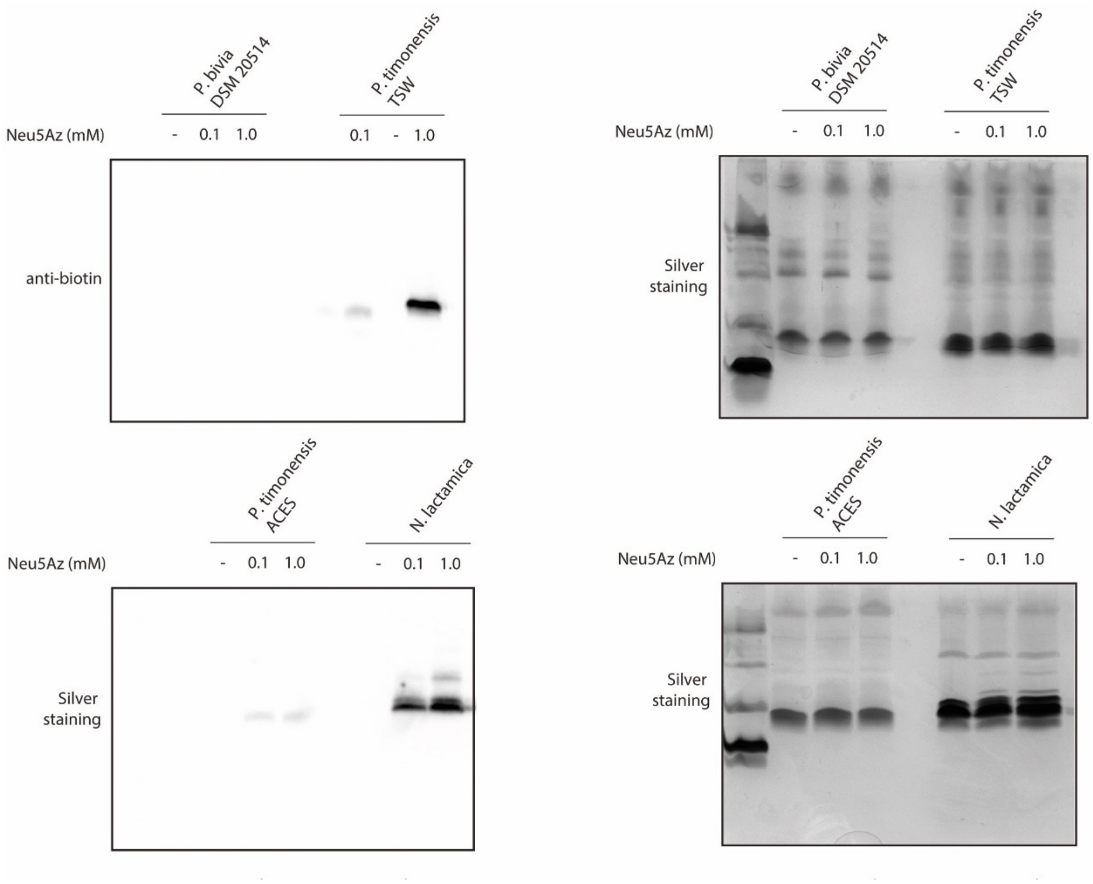


**Protein gels:**

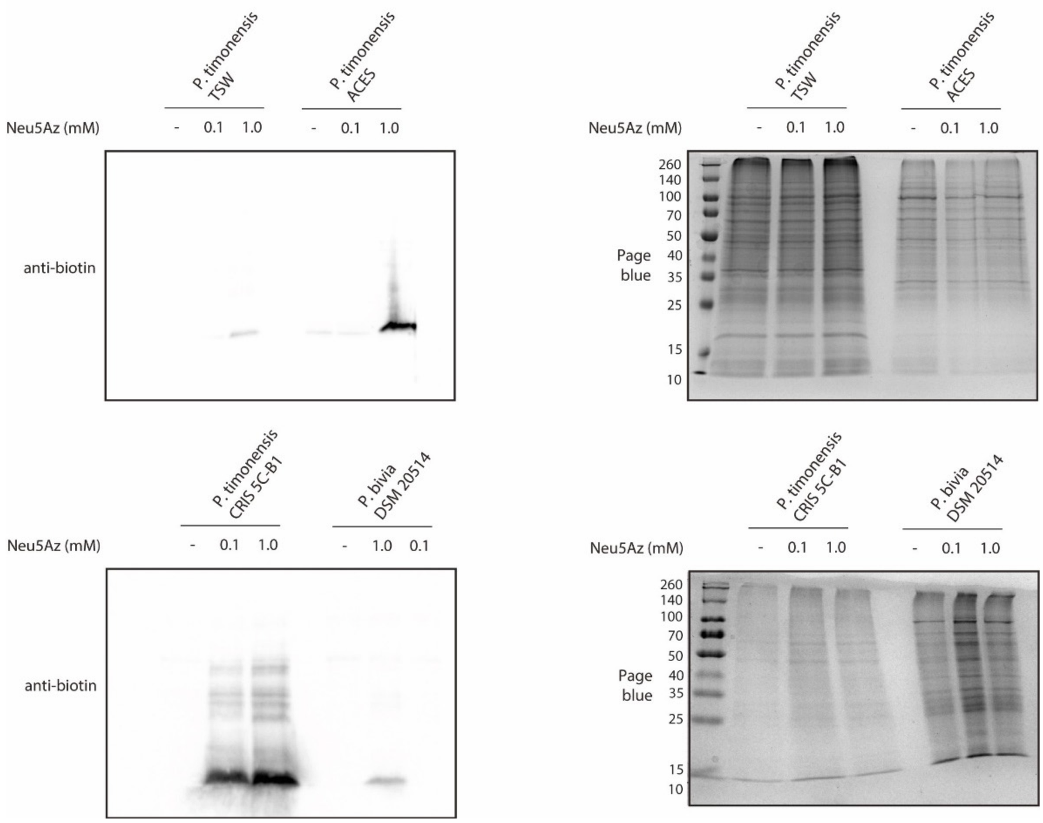

**Supplementary Figure 11.** Full length LOS gels.

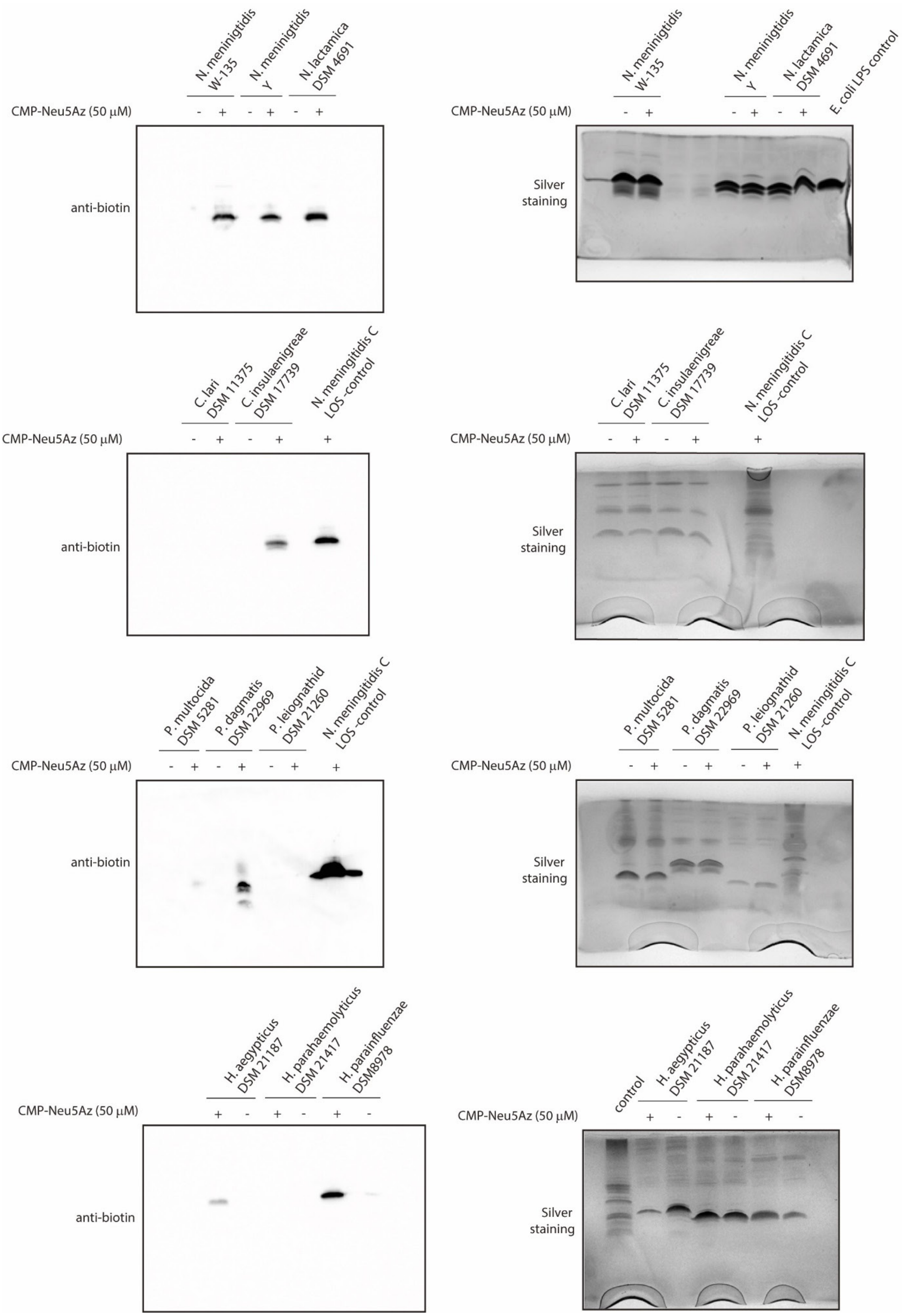

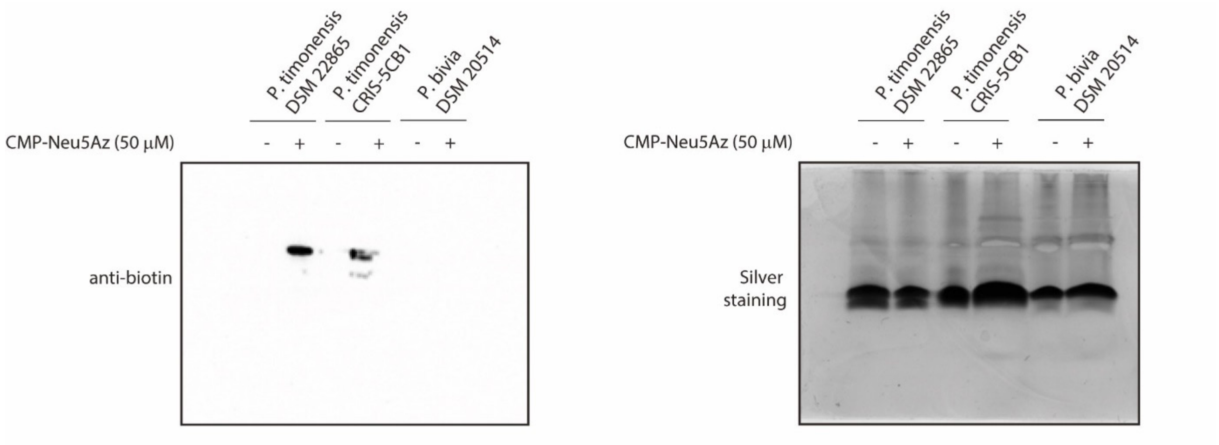
P. timonensis DSM 22865
P. timonensis CRIS-5CB1
P. bivia DSM 20514
CMP-Neu5Az (50 μM)
- + - + - +
anti-biotin
P. timonensis DSM 22865
P. timonensis CRIS-5CB1
P. bivia DSM 20514
CMP-Neu5Az (50 μM)
- + - + - +
Silver staining

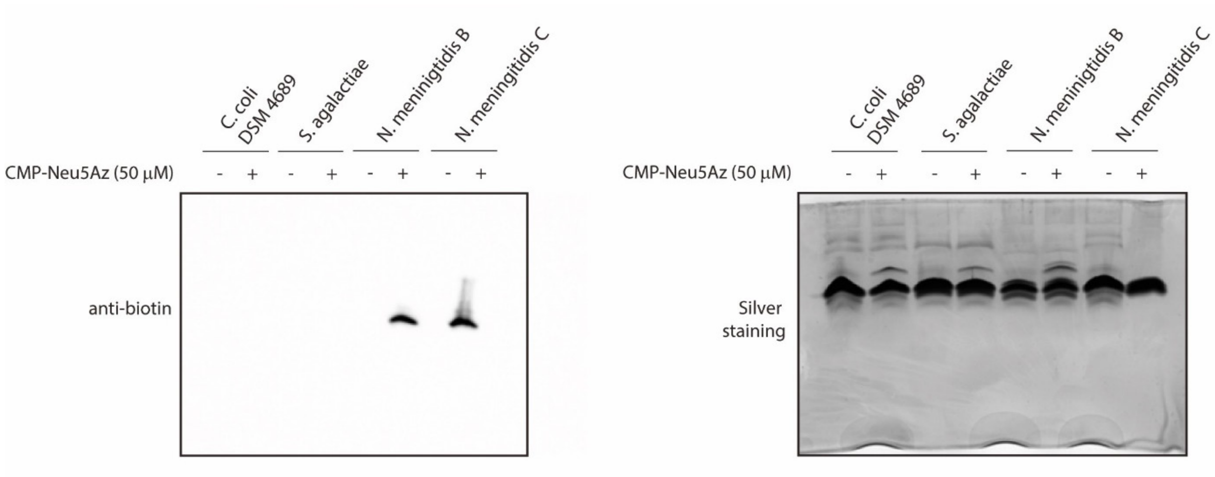
C. coli DSM 4689
S. agalactiae
N. meningitidis B
N. meningitidis C
CMP-Neu5Az (50 μM)
- + - + - + - +
anti-biotin
C. coli DSM 4689
S. agalactiae
N. meningitidis B
N. meningitidis C
CMP-Neu5Az (50 μM)
- + - + - + - +
Silver staining

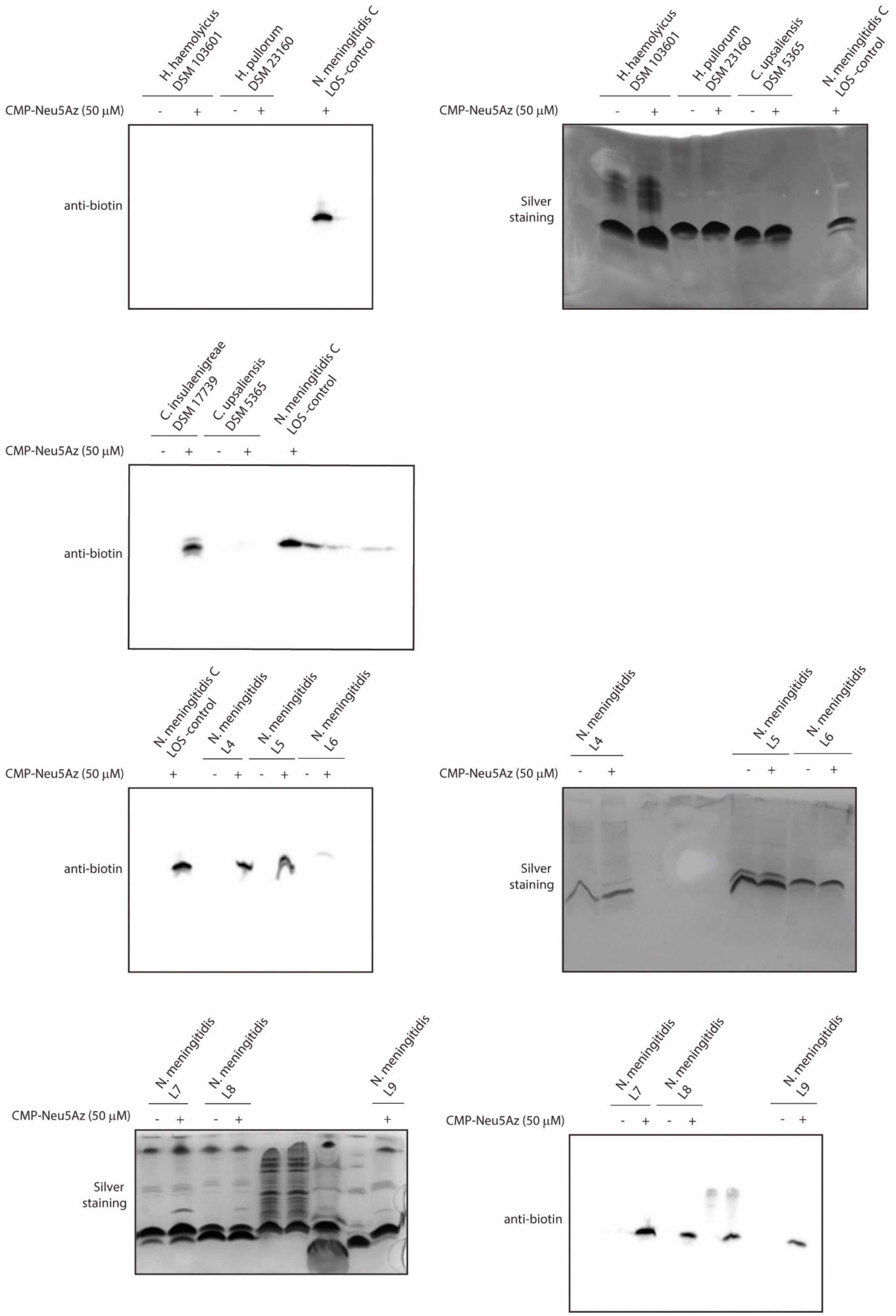
H. haemolyicus DSM 103601
H. pullorum DSM 23160
N. meningitidis C LOS-control
CMP-Neu5Az (50 μM)
- + - + +
anti-biotin
H. haemolyicus DSM 103601
H. pullorum DSM 23160
C. upsaliensis DSM 5365
N. meningitidis C LOS-control
CMP-Neu5Az (50 μM)
- + - + - + +
Silver staining
C. insulaenigreae DSM 17739
C. upsaliensis DSM 5365
N. meningitidis C LOS-control
CMP-Neu5Az (50 μM)
- + - + +
anti-biotin
N. meningitidis C LOS-control
N. meningitidis L4
N. meningitidis L5
N. meningitidis L6
CMP-Neu5Az (50 μM)
+ - + - + - +
anti-biotin
N. meningitidis L4
N. meningitidis L5
N. meningitidis L6
CMP-Neu5Az (50 μM)
- + - + - +
Silver staining
N. meningitidis L7
N. meningitidis L8
N. meningitidis L9
CMP-Neu5Az (50 μM)
- + - + +
Silver staining
N. meningitidis L7
N. meningitidis L8
N. meningitidis L9
CMP-Neu5Az (50 μM)
- + - + - +
anti-biotin

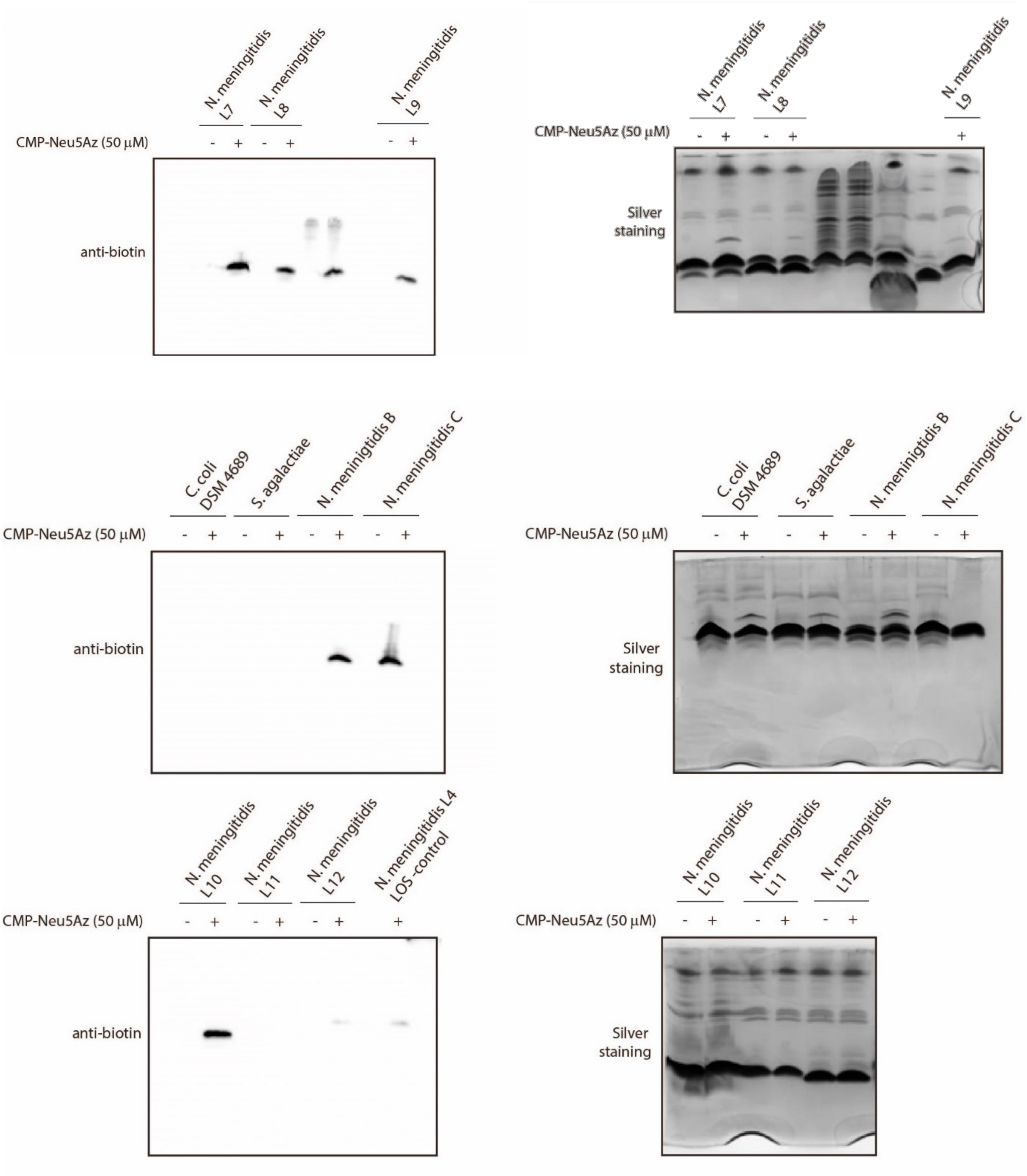
N. meningitidis L7
N. meningitidis L8
N. meningitidis L9
CMP-Neu5Az (50 μM)
anti-biotin
Silver staining
C. coli DSM 4689
S. agalactiae
N. meningitidis B
N. meningitidis C
N. meningitidis L10
N. meningitidis L11
N. meningitidis L12
N. meningitidis L4 LOS-control

**Supplementary Figure 12.** Full length gels, incubation of *N. gonorrhoeae* with CMP-Neu5Az at different time points followed by click with DBCO-biotin.

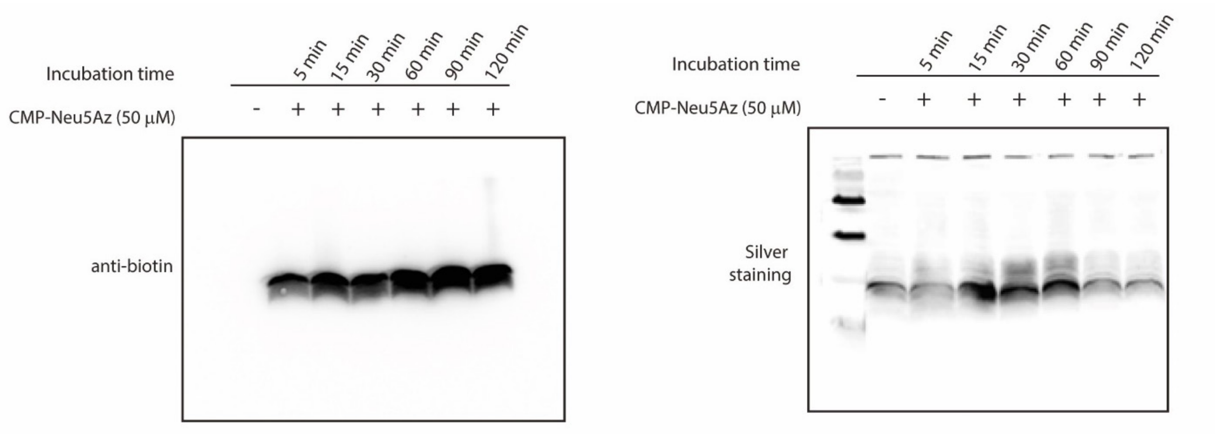


**Supplementary Figure 13.** Full length gel, **l**abelling of *C. jejuni* LOS using CMP-Neu5Az, followed by click reaction with DBCO-$AF_{488}$.

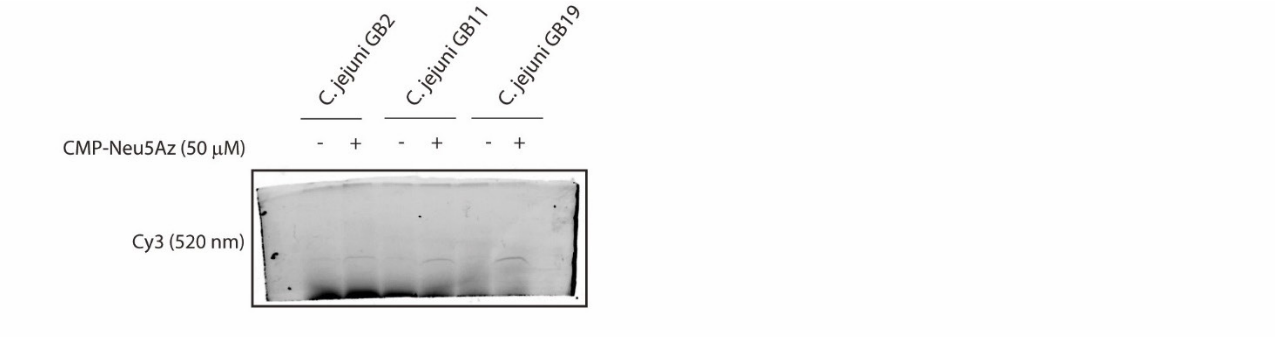
